# Myeloid-Derived Suppressor Cells are relevant factors to predict the severity of multiple sclerosis

**DOI:** 10.1101/2022.04.20.488896

**Authors:** María Cristina Ortega, Rafael Lebrón-Galán, Isabel Machín-Díaz, Michelle Naughton, Inmaculada Pérez-Molina, Jennifer García-Arocha, Jose Manuel García-Domínguez, Haydee Goicoechea-Briceño, Virginia Vila-del Sol, Víctor Quintanero-Casero, Rosa García-Montero, Victoria Galán, Celia Camacho-Toledano, María Luisa Martínez-Ginés, Denise C. Fitzgerald, Diego Clemente

**Author notes:** Corresponding author Diego Clemente, Ph.D. Grupo de Neuroinmuno-Reparación, Hospital Nacional de Parapléjicos, SESCAM, Finca “La Peraleda” s/n, 45071-Toledo, Spain. These authors contributed equally to this work (María Cristina Ortega, Rafael Lebrón-Galán).

## Abstract

Multiple Sclerosis (MS) is a highly heterogeneous demyelinating disease of the central nervous system (CNS) that needs for reliable biomarkers to foresee disease severity. Previous retrospective investigations in the MS model, experimental autoimmune encephalomyelitis (EAE), highlighted the important relationship between monocytic-myeloid-derived suppressor cells (M-MDSCs) and the experimented severity of the clinical course. In this work, we show for the first time cells resembling M-MDSCs associated to MS lesions, whose abundance was related to milder MS clinical courses. Moreover, Ly-6C^hi^ cells (which are indistinguishable from circulating M-MDSCs in mice) are useful biomarkers to predict a milder severity of the EAE disease course and a lesser tissue damage extent. Finally, the abundance of M-MDSCs in blood from untreated MS patients at their first relapse was inversely correlated with EDSS at baseline and relapse recovery one-year later. In summary, our data point to M-MDSC load as a promising biomarker of patient’s clinical course severity.

**Teaser:** The abundance of myeloid-derived suppressor cells is related to a milder clinical course in multiple sclerosis patients.

## INTRODUCTION

Multiple sclerosis (MS) is a chronic, immune-mediated and demyelinating disease of the central nervous system (CNS). Following traumatic lesions, it is the main driver of CNS disability in young adults (*1*). MS remains incurable, and its course and response to treatments vary considerably among patients (*2*). Around 85% of patients present the relapsing-remitting clinical form of MS (RRMS), which is characterized by an initial episode of neurologic dysfunction, followed by subsequent periods of remission (clinical recovery) and relapsing disease (*3*). This pattern of disease has led to the belief that molecules or cell types that control the immune system activation play a relevant role in MS (*4*). In this regard, approved disease-modifying treatments (DMTs), mainly effective in relapsing forms of MS, had helped shed light on the crucial role of immunoregulatory mechanisms in the recovery from MS relapses (*5*).

The wide variability in the MS clinical course severity poses a challenge to neurologists when choosing among DMTs. Although magnetic resonance imaging (MRI) allows us to obtain clinical and diagnostic information, and helps monitoring the treatments (*1, 6*), no single biochemical/immune biomarker can currently predict disease severity in each patient. More intense inflammation, as a result of the increase of pro-inflammatory cytokines detected in the cerebrospinal fluid (CSF) and the meninges of MS samples, is associated with a higher degree of cortical demyelination, which is related to a more severe clinical course (*7*). Notably, patients with a severe clinical course have a higher lesion load and more interestingly, a higher proportion of active lesions (*8*). However, little is known about the relationship between regulatory myeloid cells and the severity of the clinical course of MS (*9, 10*). Therefore, information about the alterations to the number and/or activity of myeloid regulatory cells in MS patients, and the consequences of these, may help us elucidate some of the less well understood issues regarding MS, such as the highly heterogeneous clinical course of the disease.

Myeloid-derived suppressor cells (MDSCs) have gained importance as pivotal factors involved in the regulation of immune response, generating particular interest in cancer and immune-mediated diseases like MS (*11*). MDSCs comprise two main subsets of suppressive immune cells: polymorphonuclear-MDSCs (PMN-MDSCs) and monocytic-MDSCs (M-MDSCs), which are characterized by the differential expression of specific Gr-1 epitopes (Ly-6C and Ly-6G) in murine cells and of CD14/CD15 markers in humans (*11*). The immunoregulatory role of MDSCs has been demonstrated in the experimental autoimmune encephalomyelitis (EAE) animal model of MS (*12, 13*), where M-MDSCs (CD11b^+^ Ly-6C^hi^ Ly-6G^-/low^, which are phenotypically indistinguishable from Ly-6C^hi^ inflammatory monocytes, i.e. Ly-6C^hi^ cells (*14*), have a stronger immunosuppressive activity than PMN-MDSCs (CD11b^+^ Ly-6C^int^ Ly-6G^hi^) (*12, 15*). The presence of Arg-I^+^ cells has been identified in the CNS of mice with EAE, showing typical surface markers for M-MDSCs (*13*). Furthermore, there is a correlation between the abundance of Ly-6C^hi^ cells in the spleen and the previous severity of the clinical disease course in EAE (*13, 16*). Importantly, pharmacological compounds can improve the immunosuppressive activity of Ly-6C^hi^ cells in the induction phase of an animal model of primary progressive MS (PPMS), as well as in the EAE model (*17, 18*). As such, Ly-6C^hi^ cells/M-MDSCs are promising targets for potential therapies in autoimmune diseases.

The identification of this cell population in human diseases has recently reached an agreement on its phenotypic classification, being M-MDSCs classified as CD11b^+^ CD33^+^ HLA-DR^-/low^ CD14^+^ CD15^-^ cells, while the PMN-MDSC subset are CD11b^+^ CD33^+^ HLA-DR^-/low^ CD14^-^ CD15^+^ LOX1^+^ cells (*11, 19, 20*). Currently, MDSCs have been mainly found in the peripheral blood of MS patients, albeit not without some controversy. Some authors described an increase in the abundance of PMN-MDSCs in MS patients during remission (*20*), while others observed more PMN-MDSCs and M-MDSCs during relapses (*21, 22*). Notably, M-MDSCs show a stronger immunosuppressive function during relapses than remission and the proportion of M-MDSCs in the peripheral blood of MS patients is 10-fold higher than that of PMN-MDSCs (*20, 22*). Hence, M-MDSCs appear to be the most relevant subset to analyse when searching for therapeutic strategies that might enhance immune-regulation during MS relapse. However, there are no data about the relationship of M-MDSCs and the clinical severity of MS, or about their presence in the CNS of MS patients, important prior steps before considering their possible use as a decision-making tool for neurologists.

In the present paper, we describe for the first time that infiltrated myeloid cells showing the phenotypic markers of M-MDSCs are present in tissue from progressive MS patients with different disease duration, and we show similar results in both the murine model and human pathology. We describe that the abundance of Ly-6C^hi^ cells in peripheral blood at the onset of EAE is inversely correlated with the severity of the future clinical course and the degree of histopathological damage in the spinal cord. Furthermore, our data from untreated MS patients show an inverse correlation between the abundance of M-MDSCs measured at an early time point of their clinical course and the EDSS at baseline and one year later. Hence, the data presented here suggest that peripheral M-MDSCs could be used as biomarker for MS prognosis to predict the severity of the clinical course in MS.

## RESULTS

### M-MDSCs are present in the CNS of MS patients

Although differences in the abundance of M-MDSCs in the peripheral blood of MS patients have been reported (*20–23*), these cells have yet to be described in MS tissue. We explore the presence of cells with the M-MDSC phenotype in human tissue by using the commonly markers for their identification in the human peripheral blood, i.e. CD14^+^ CD15^-^ CD11b^+^ HLA-DR^-/low^. CD14 and CD15 are pivotal markers to classify human monocyte or polymorphonuclear subpopulations (*11*), so we established the distribution pattern of both markers in MS lesions. While in control human tissue neither CD14 nor CD15 were observed in the normal white matter (Fig. 1A-C), CD14^+^ cells were observed mainly within the demyelinated plaque of active lesions (AL) and in the rim of mixed active/inactive lesions (rAIL; Fig. 1D-E, G-H), whereas they were virtually absent from the center of mixed active/inactive lesion (cAIL) and inactive lesions (IL; Fig. 1G-H, J-K). By contrast, no CD15 was detected in any MS lesion (Fig. 1F, I, L), but was rather restricted to perivascular granulocytic-like cells (Fig. 1M-O).

**Figure 1:**
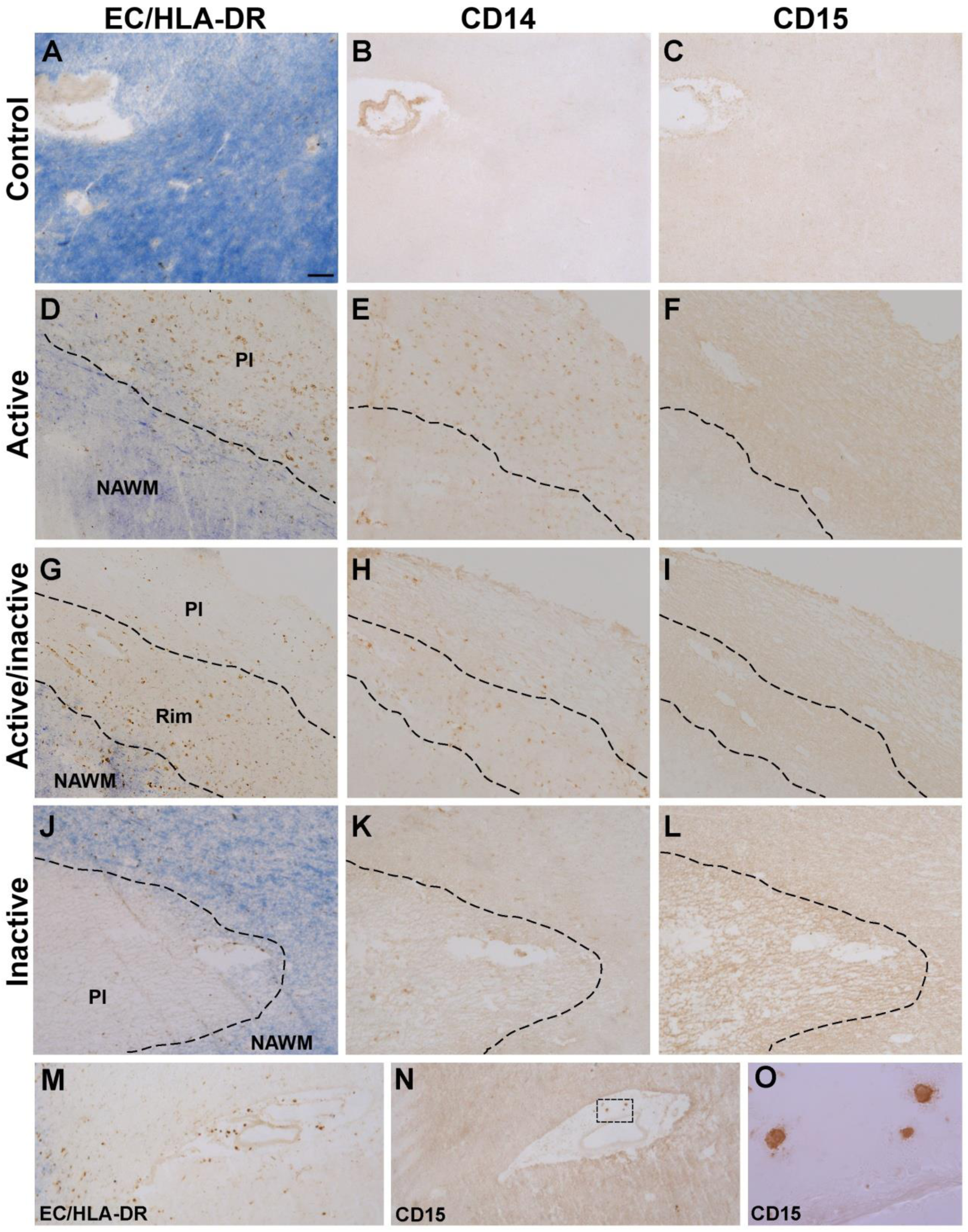
Expression pattern of CD14 and CD15 in MS lesions. **A-C:** CD14 or CD15 staining was not detected in control human tissue. **D-O:** CD14^+^ cells were located both in the plaque of AL (**D**, **E**) and in the rim of AIL (**G, H**). By contrast, these cells were almost absent in the center or plaque of AIL (**G**, **H**) and IL (**J, K**). CD15 staining was not detected in any region of the MS lesions (**F**, **I**, **L**), although CD15^+^ granulocytes were clearly detected in the perivascular area of blood vessels (**M**-**O**). EC, eriochrome cyanine; AL, active lesions; AIL, mixed active/inactive lesions; IL, inactive lesions; Pl, plaque; NAWM, normal appearing white matter. Scale bar: A-C, J-N = 100 µm; D-I = 125 µm; O = 15 µm.

HLA-DR, CD15 and CD11b staining was carried out to complete the characterization of M-MDSCs. CD14^+^ cells in MS lesions expressed varying intensities of HLA-DR staining (Fig. 2A-C). According to previous classifications of M-MDSCs based on the intensity of HLA-DR staining (*11*), we established two cell subpopulations, whereby CD14^+^ HLA-DR^-/low^ cells were considered as M-MDSC-like cells (Fig. 2A-B) and CD14^+^ HLA-DR^hi^ cells were identified as putative inflammatory macrophages (Fig. 2C). Moreover, we corroborated the classification of these cells as putative M-MDSCs through the lack of CD15 staining in the CD14^+^ cell population (Fig. 2A-C). Finally, all CD14^+^ cells in MS lesions expressed the myeloid cell marker CD11b (Fig. 2D-G), which was in agreement with their myeloid cell nature. In order to check the infiltrating origin of the M-MDSC-like cells, we labeled with CD14, HLA-DR and TMEM119, as a specific marker for microglia in human tissue able to discriminate these myeloid resident cells from monocyte-derived infiltrated macrophages (*24–26*). First we checked whether TMEM119^+^ cells far from MS lesions showed the morphology of microglial cells (Fig. 2H-K). Within MS demyelinating lesions, less brilliant TMEM119 immunoreactivity was also present in round-shaped amoeboid/globular cells (Fig. 2L-O). In this sense, CD14^+^HLA-DR^hi^ cells resembling inflammatory macrophages were observed positive or negative for TMEM119, pointing to their microglial or monocyte-derived origin, respectively. Interestingly, CD14^+^HLA-DR^-/low^ M-MDSC-like cells always lacked TMEM119 immunoreactivity (Fig. 2L-O), in agreement with their peripheral origin as monocyte-derived cells. In summary, infiltrated cells with the M-MDSC phenotype were characterized as CD14^+^ HLA-DR^-/low^CD15^-^ CD11b^+^ TMEM119^-^, showing for the first time, the presence of myeloid cells with typical markers for M-MDSCs (hereinafter referred to as CD14^+^HLA-DR^-/low^ M-MDSCs) within the CNS of MS patients.

**Figure 2:**
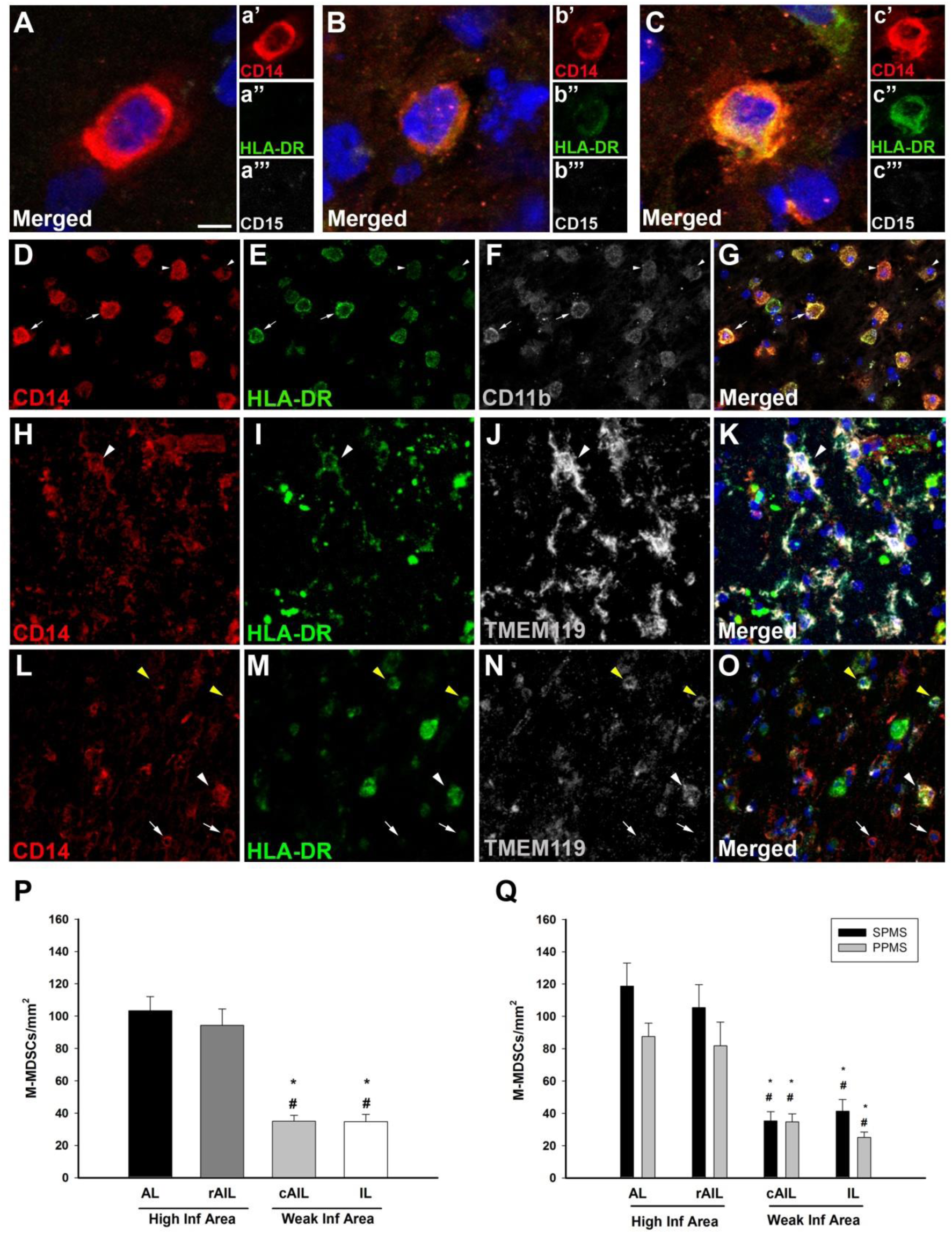
Cells resembling infiltrated M-MDSC are present within MS lesions. **A-C:** CD14^+^ cells showed different levels of HLA-DR in demyelinating lesions. Based on the intensity of HLA-DR (á′, b′′, ć′) we identified two cell subpopulations: CD14^+^ HLA-DR^-/low^ cells were considered as M-MDSC-like cells (**A**, **B**) and CD14^+^ HLA-DR^hi^ cells were identified as putative inflammatory macrophages (**C**). The lack of CD15 expression in CD14^+^ cells (á′′, b′′′, ć′′) corroborated the exclusive presence of M-MDSC-like cells in MS lesions. **D**-**G**: Both CD14^+^ cell subpopulations were stained for CD11b, confirming their myeloid cell nature. Arrowheads point to myeloid cells with M-MDSC phenotype, i.e CD14^+^CD11b^+^HLA-DR^low^, while arrows indicate CD11b^+^ inflammatory macrophages. **H**-**O**: TMEM119^+^ cells with microglial morphology were detected in the NAWM (**H**-**K**). The lack of TMEM119 immunoreactivity observed in amoeboid CD14^+^HLA-DR^-/low^ M-MDSC-like cells (pointed by arrows on L-O) confirmed the peripheral origin of these cells. Yellow and white arrowheads in L-O point to inflammatory monocyte-derived macrophages (CD14^+^ HLA-DR^hi^) and microglial-derived macrophages (CD14^-^ HLA-DR^hi^), respectively. **P**: Quantification of the density of putative M-MDSC in all MS lesion types from SPMS and PPMS patients. The abundance of CD14^+^ CD15^-^ HLA-DR^-/low^ cells was significantly higher in the AL and in the rAIL (n=46 AL, 27 AIL and 24 IL from 33 MS patients). **Q:** AL and the rAIL from SPMS and PPMS patients showing a greater presence of M-MDSC-like cells than in cAIL and IL (n = 24 AL, 14 AIL and 14 IL from 20 SPMS patients; 22AL, 12 AIL and 10 IL from 13 PPMS patients). Scale bar: A-C = 5 µm; insets in A-C = 10 µm; D-G = 40 µm; H-K = 20 µm; L-O = 37 µm. Comparative analyses were carried out with ANOVA and the mean + SEM data are shown: *, # p < 0.05. In Q, * represents the difference with respect to AL and # with respect to the rAIL. rAIL, rim of AIL; cAIL, center of AIL; High. Inf Area, high inflammatory area; Weak Inf Area, weak inflammatory area.

Once characterized, we quantified the M-MDSC density in the different types of demyelinated lesions. These cells were mainly distributed in the demyelinating area of AL and in the rAIL, where their density was significantly higher than in the cAIL and IL (Fig. 2P). Given that these regions are characterized by different inflammatory activity and the spontaneous capacity for remyelination (*27*), we gathered the data obtained from the AL and the rAIL (which from here on will together be referred to as the high inflammatory area), and compared this with the results from the cAIL and IL (hereafter referred to as the weak inflammatory area). As a second step, we analyzed the distribution of these cells based on the classification of the patient’s clinical course, secondary progressive MS (SPMS) and primary progressive MS (PPMS). In this sense, we found there was a similar M-MDSC distribution in all regions analyzed from SPMS and PPMS patients, with the highest density of M-MDSCs in high inflammatory areas (Fig. 2Q; Fig. S2A).

In summary, we provide the first description of a myeloid cell population expressing typical markers to be considered as infiltrated M-MDSCs in the CNS from MS patients and report that these cells are found mainly in areas of high inflammatory activity. Furthermore, there appears to be no difference in the distribution of M-MDSCs in the CNS of SPMS and PPMS patients.

### M-MDSCs are differentially associated with the severity of the clinical course in SPMS and PPMS patients

In order to explore the influence of disease severity on M-MDSC distribution, we compared their density in the CNS of patients with a short or long clinical course according to the median value of the patients’ disease duration from the SPMS or PPMS sample collection (Table 1).

**Table 1:** Demographic and clinical characteristics of MS patients and control subjects. CNS samples from MS patients provided by the UK MS Tissue Bank (London, UK, identified as MS) and Dame Ingrid V. Allen tissue collection (from Belfast, UK, identified as MSD). AL: active lesion; AIL: mixed active/inactive lesion; Ct: control; DD: disease duration; F: female; h: hours; IL: inactive lesion; M: male; nd: Not determined; PP: primary progressive; SP: secondary progressive; TP: time *post-mortem,* y: years.

| Patient | Clinical course | MS | Age/Sex | TP (h) | DD (y) | Type of lesion |  |  |
| --- | --- | --- | --- | --- | --- | --- | --- | --- |
|  |  |  |  |  |  | AL | AIL | IL |
| MS122 | Short<br>(DD ≤ 16 y) | SP | 44/M | 16 | 16 |  |  | 1 |
| MS356 |  | SP | 45/F | 10 | 16 | 1 |  |  |
| MS408 |  | SP | 39/M | 21 | 10 | 2 | 2 |  |
| MS422 |  | SP | 58/M | 25 | 13 | 1 |  | 1 |
| MS371 |  | SP | 40/M | 27 | 14 | 1 |  |  |
| MS114 |  | SP | 52/F | 12 | 15 |  |  | 1 |
| MS163 |  | SP | 45/F | 28 | 6 | 1 |  |  |
| MS 154 |  | SP | 34/F | 12 | 12 | 1 | 1 | 1 |
| MS160 |  | SP | 44/F | 18 | 16 | 4 | 1 | 1 |
| MSD15 |  | SP | 38/M | nd | 14 | 1 |  |  |
| MSD70 |  | SP | 49/M | nd | 13 |  | 2 |  |
| MS318 | Long<br>(DD > 16 y) | SP | 59/F | 13 | 34 | 3 | 2 |  |
| MS187 |  | SP | 57/F | 13 | 27 |  | 1 | 1 |
| MS404 |  | SP | 55/F | 17 | 34 | 1 | 2 | 1 |
| MS466 |  | SP | 65/F | 25 | 36 | 1 | 1 |  |
| MS406 |  | SP | 62/M | 23 | 47 | 3 |  |  |
| MS470 |  | SP | 64/F | nd | 35 | 2 | 2 | 2 |
| MS055 |  | SP | 47/F | 15 | 32 | 2 | 1 | 3 |
| MS330 |  | SP | 59/F | 21 | 39 |  |  | 1 |
| MSD58 |  | SP | 82/M | nd | 62 |  |  | 1 |
| MS383 | Short<br>(DD ≤ 17 y) | PP | 42/M | 17 | 17 | 3 | 1 |  |
| MS325 |  | PP | 51/M | 13 | 2 | 1 |  |  |
| MS083 |  | PP | 54/M | 13 | 13 |  | 1 | 2 |
| MS094 |  | PP | 42/F | 11 | 6 | 3 | 3 | 1 |
| MS473 |  | PP | 39/F | 9 | 13 | 4 | 2 |  |
| MS216 |  | PP | 58/F | 9 | 12 | 1 |  |  |
| MS500 |  | PP | 50/M | 32 | 8 | 5 |  |  |
| MSD276 |  | PP | 40/F | nd | 17 |  | 1 |  |
| MS057 | Long<br>(DD > 17 y) | PP | 77/F | 9 | 47 | 1 | 1 | 1 |
| MS168 |  | PP | 88/F | 22 | 30 | 2 |  | 2 |
| MS492 |  | PP | 66/F | 15 | 31 |  |  | 1 |
| MS398 |  | PP | 57/F | 28 | 28 |  |  | 1 |
| MS363 |  | PP | 42/M | 20 | 27 | 2 | 3 | 1 |
| CO22 |  | Ct | 69/F | 33 |  |  |  |  |
| CO72 |  | Ct | 72/M | 26 |  |  |  |  |
| CO73 |  | Ct | 71/M | 29 |  |  |  |  |
| CO74 |  | Ct | 84/F | 22 |  |  |  |  |
| PDCO23 |  | Ct | 78/F | 23 |  |  |  |  |
| PDCO28 |  | Ct | 84/F | 11 |  |  |  |  |

We did not observe differences between SPMS patients with a long (Fig. 3A-F) or short disease duration (Fig. 3G-L), being the highest M-MDSC density always present in areas of high inflammation (Fig. 3A-C, G-I, M). In order to scrutinize whether the balance between pro-inflammatory/immunossuppresive cells would be affected by the disease course, the ratio of CD14^+^ HLA-DR^hi^ cells/M-MDSCs (hereinafter referred to as inflammatory/immunosuppressive ratio-IIR) was calculated in MS lesions. Similar to M-MDSCs, the IIR was always significantly higher in high inflammatory regions of both SPMS and PPMS (Fig. S2B). On the other hand, this ratio was increased in high inflammatory areas from SPMS patients with short or long clinical course, irrespectively of the disease duration (Fig. 3N). Interestingly, M-MDSCs were not equally distributed in PPMS patients. The distribution of these cells in PPMS with a long disease (Fig. 4A-F, M) was similar to that in SPMS patients (Fig. 3M), being always the density of M-MDSC significantly higher in high inflammatory regions. On the contrary, PPMS patients with a short clinical course showed no differences in the M-MDSC density between high and weak inflammatory areas (Fig. 4G-L, M). Interestingly, the density of this regulatory cell population in regions with a high inflammatory activity was significantly lower in those PPMS patients with a short disease course (Fig. 4M). In addition, IIR was dramatically increased in PPMS with short disease durations, suggesting a dampened immunoregulatory environment (Fig. 4N).

**Figure 3:**
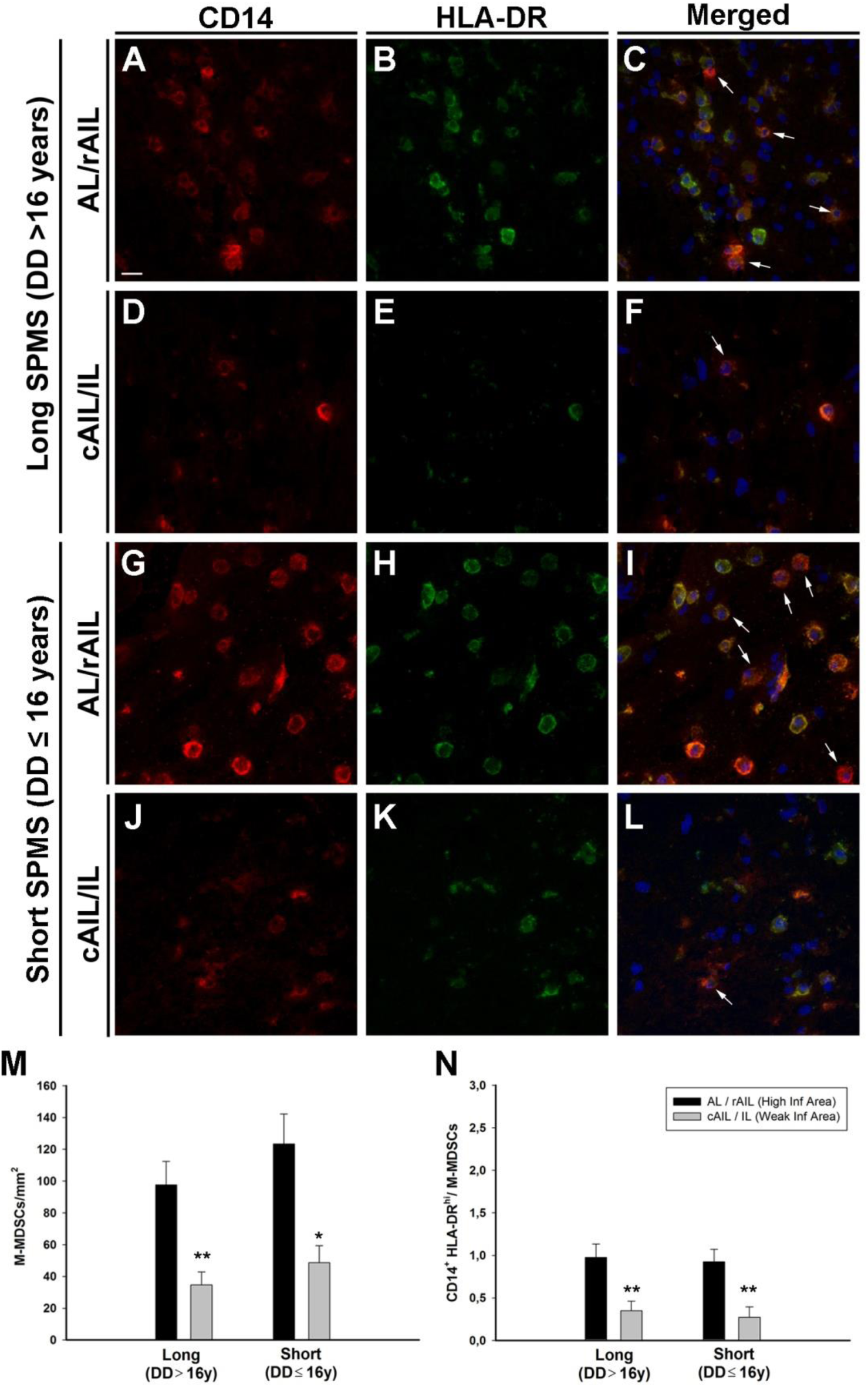
Distribution pattern of M-MDSCs in SPMS patients with different clinical course attending to the disease duration. **A-L:** The highest density of M-MDSC was always seen in high inflammatory areas, independent of the disease duration (long DD in **A**-**C**, short DD in **G**-**I**). Both the cAIL and IL from SPMS with a long (**D**-**F**) and short disease duration (**J**-**L**) have a low density of M-MDSC. The arrows indicate M-MDSC (CD14^+^ CD15^-^ HLA-DR^-/low^). **M**: Quantification of the M-MDSC distribution showing that the disease duration does not affect their density, which was always significantly higher in high inflammatory areas. N: Ratio between CD14^+^ HLA-DR^hi^ cells/M-MDSCs (also referred as IIR in the main text) was significantly increased in high inflammatory areas of SPMS, independently on the disease duration. N = 11 SPMS with short DD and 9 SPMS with long DD. Scale bar: A-L = 20 µm. A comparative analysis was carried out with a Student’s *t* test, the means (± SEM) data are indicated: *p < 0.05, **p < 0.01 and ***p < 0.001. DD, disease duration. IIR: inflammatory-immunoregulatory ratio.

**Figure 4:**
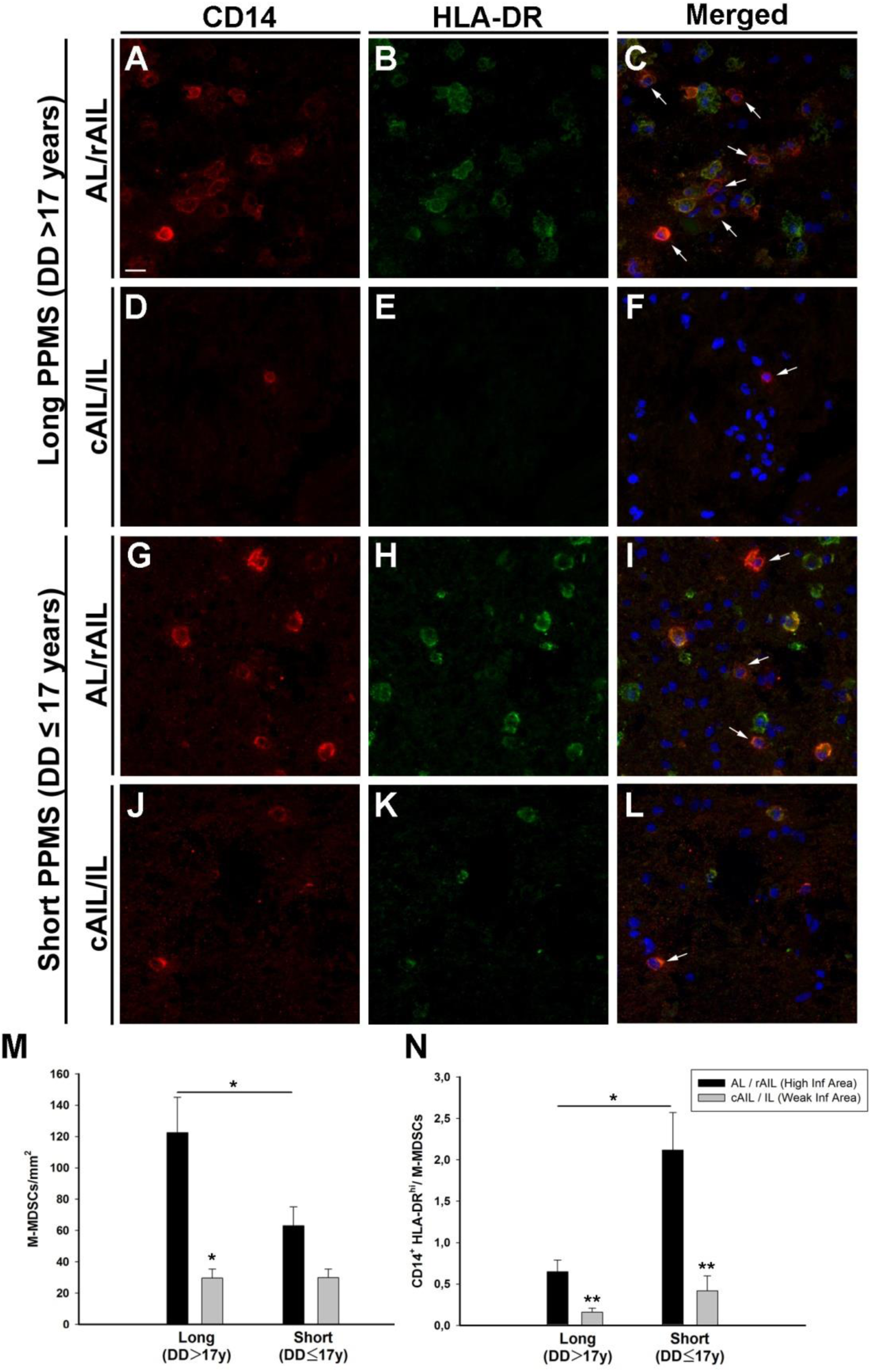
Disease duration affects the M-MDSC distribution in PPMS patients. **A-L:** In the high inflammatory area (**A-C**) there was a higher density of M-MDSCs than in the weak inflammatory regions (**D-F**) of PPMS patients with long disease duration. By contrast, the density of this cell population in both regions was similar in the PPMS patients with a short clinical course (**G-L**). **M**: Only PPMS patients with a long progressive course had a significantly higher abundance of M-MDSCs in the high inflammatory areas, while the distribution of these cells in the PPMS patients with short disease duration was similar between the intense and weak inflammatory areas. N: the IIR significantly increased in higher inflammatory regions in those PPMS patients with a short disease duration (n = 8 PPMS with a short DD and 5 PPMS with a long DD). Scale bar: A-L = 20 µm. A comparative analysis was carried out with a Student’s *t* test and the mean (± SEM) data are shown: *p < 0.05 and **p < 0.01. IIR: inflammatory-immunoregulatory ratio.

Based on these results, we also analyzed the correlation between M-MDSC density or the IIR in MS lesions and the disease duration. In the SPMS, M-MDSCs density or the IIR were independent of the disease duration in both high (r = 0.154, p = 0.583) and weak inflammatory areas (r = 0.081, p = 0.773; Fig. 5A-B). Similarly, both parameters were independent of the time *post-mortem* or the patient’s age (Table S1). In parallel, the presence of M-MDSCs in high inflammatory regions of PPMS was not correlated with time *post-mortem* (Table S1). Interestingly, the abundance of M-MDSCs in high inflammatory areas of PPMS patients was directly correlated with age and disease duration, i.e. the lower the M-MDSC density in regions with a high inflammatory activity, the younger and the shorter clinical course duration (Fig. 5C; Table S1). In addition, the disease duration in PPMS patients showed an inverse correlation with the IIR in high inflammatory regions, suggesting that the more dampened inmunoregulatory environment, the shorter disease duration (Fig. 5D). Finally, no correlations were evident between the M-MDSC density or the IIR in weak inflammatory areas and the disease duration (r = 0.299, p = 0.384 and r = -0.123, p = 0.693, respectively), the age or the time *post-mortem* in PPMS patients (Table S1).

**Figure 5:**
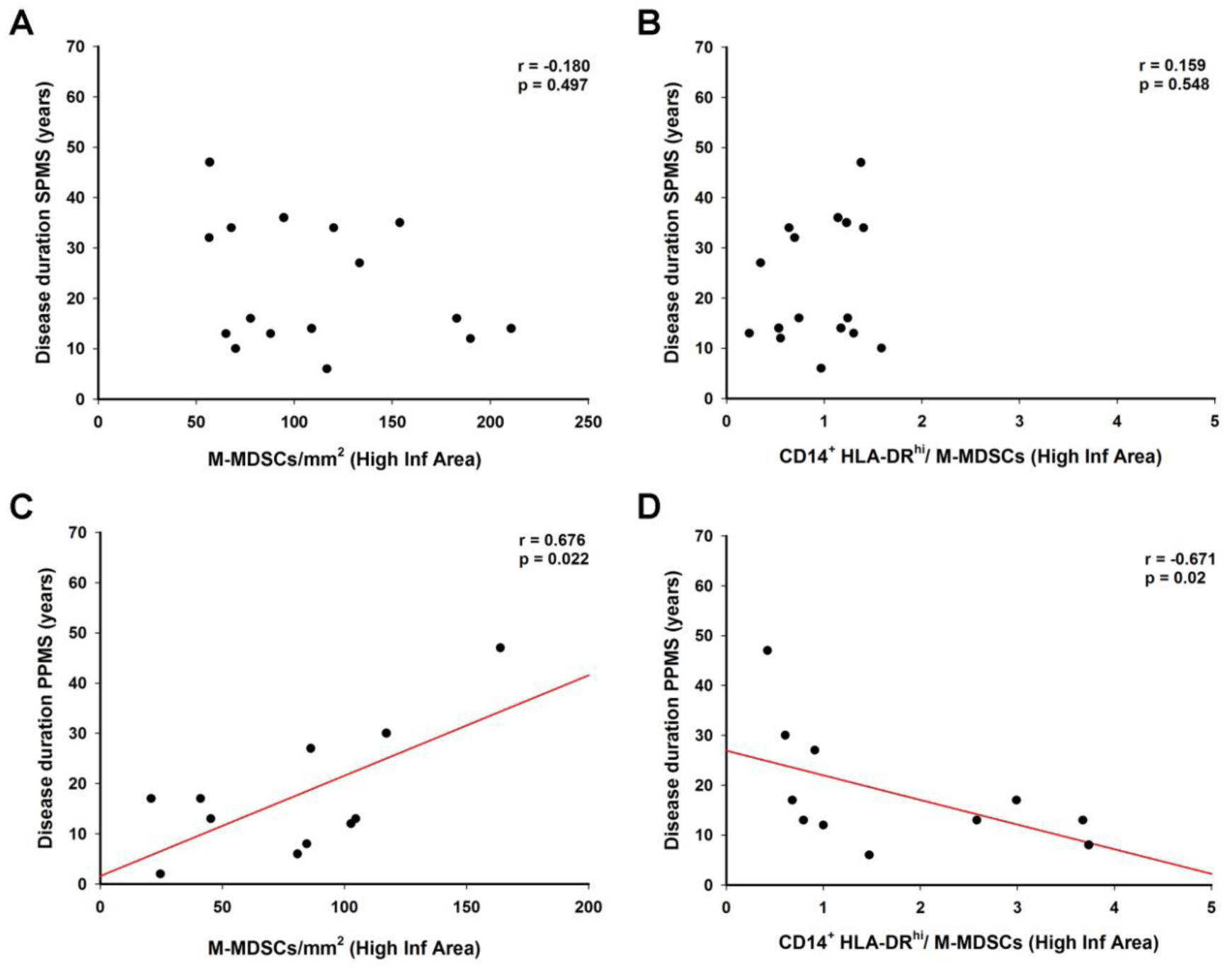
Correlation analysis of the M-MDSC density and the inflammatory cells/M-MDSCs ratio with the disease duration of MS patients. **A-D:** In the high inflammatory regions, there was no correlation between the density of M-MDSCs (**A**) or the IIR (**B**) and the disease duration in SPMS patients (n=16 SPMS in A and n= 14 SPMS in B). The abundance of this regulatory cell population in high inflammatory areas from PPMS patients was directly correlated with the duration of the clinical course (**C**), while this correlation was not seen in the weak inflammatory areas (**D**; n=11 PPMS in C and n= 10 PPMS in D). For the correlation analysis, a Spearman test was carried out. IIR: inflammatory-immunoregulatory ratio.

In summary, these findings demonstrate that the distribution of myeloid cells expressing typical markers for M-MDSCs is independent to clinical course duration in SPMS patients. However, a lower density of these regulatory cells in parallel with an exacerbated inflammatory context is directly correlated with a shorter disease durations in PPMS patients. However, a main open question is whether M-MDSCs may be considered a valuable tool for predicting the severity of the clinical course in MS.

### A higher abundance of blood Ly-6C^hi^ cells at onset is indicative of a milder EAE disease severity

As a first step to establish the value of M-MDSCs to predict the future disease progression in MS, we explore the relationship of their abundance in the blood and the future clinical sign dimension in EAE mice. M-MDSCs with a high immunosuppressive activity have been phenotipically characterized in the spleen and the spinal cord of EAE mice as CD11b^+^ Ly-6C^hi^ Ly-6G^-/low^, also called Ly-6C^hi^ cells (*12, 28*). However, it has been repeatedly demonstrated that circulating murine M-MDSCs are phenotipically indistinguishable from Ly-6C^hi^ inflammatory monocytes in different inflammatory contexts (*12, 14, 19, 28, 29*). Our previous data established a clear relationship between splenic M-MDSCs (i.e. Ly-6C^hi^ cells) at the peak of the clinical course and the previous experimented severity of EAE mice (*16*). For all these reasons, in order to validate M-MDSCs as a predictive tool for MS severity, we firstly explored the correlation between the abundance of cells sharing their immunophenotype, i.e. Ly-6C^hi^ cells, in the peripheral blood at disease onset and different clinical parameters during the EAE clinical course, including the severity index (SI) (*16*). We showed that the higher abundance of Ly-6C^hi^ cells was related to a less aggressive clinical course according to the SI (r = -0.888, p<0.000001: Fig. 6A-B). Similar, but more modest correlations were observed with the maximum (Fig. 6C) and the accumulated clinical score (Fig. 6D). In our hands, 20% of mice that were immunized did not develop clinical signs of EAE. Interestingly, when the peripheral blood of mice with EAE was collected before clinical onset (at 10 dpi), there were significantly fewer Ly-6C^hi^ cells in mice that went on to develop EAE than in those that remained asymptomatic (Fig. 6E).

**Figure 6:**
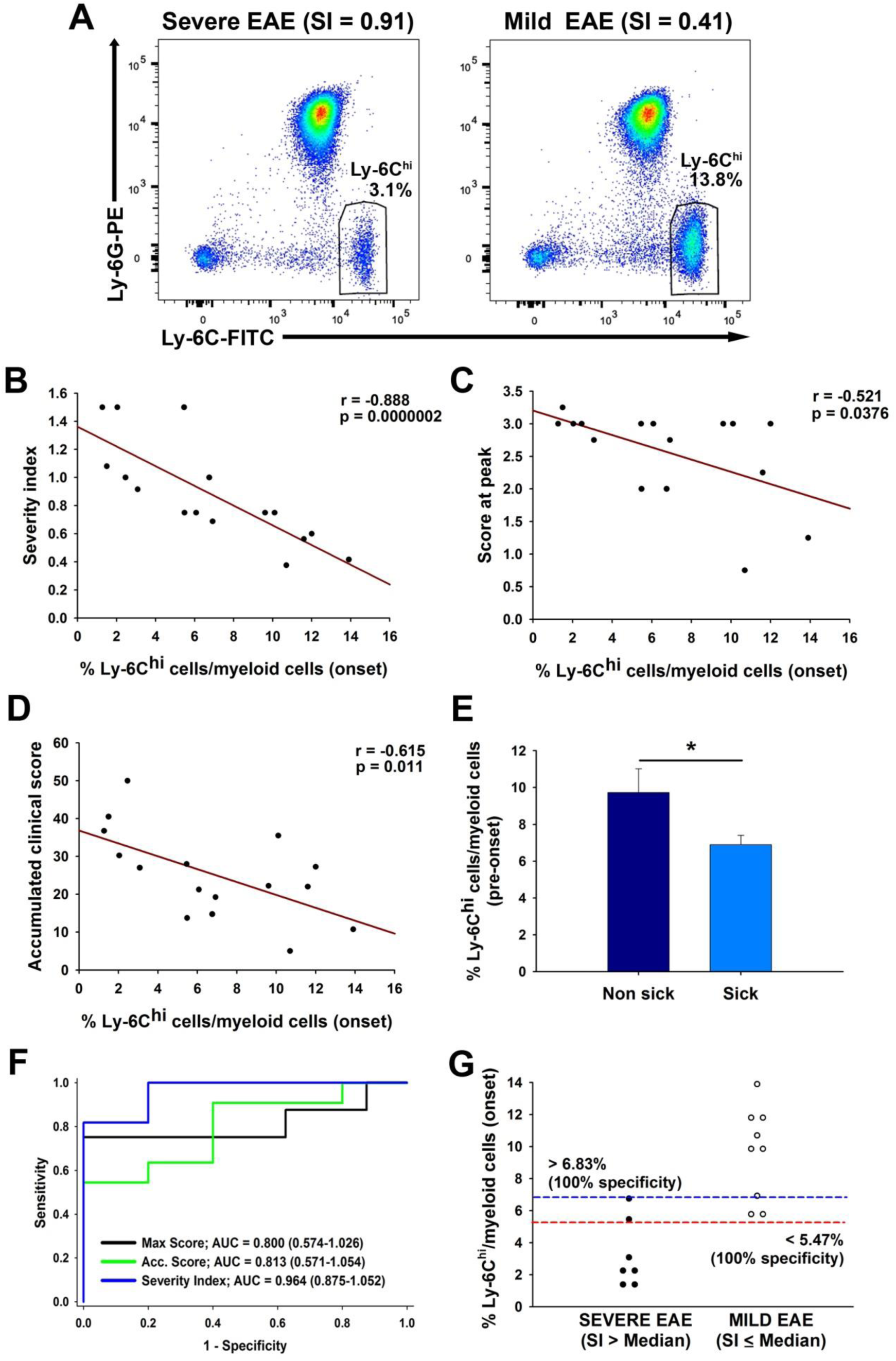
The proportion of Ly-6C^hi^ cells in the peripheral blood is indicative of a less severe clinical evolution. **A**: Representative flow cytometry plots of Ly-6C^hi^ cells from EAE mice with a severe (left panel) or mild (right panel) clinical course. **B-D**: The abundance of Ly-6C^hi^ cells relative to the myeloid component at the onset of the clinical symptoms was inversely correlated with the severity index (SI, **B**) of the disease course. The maximal (**C**) and the accumulated clinical score (**D**) were also inversely correlated with the amount of Ly-6C^hi^ cells, yet in a milder manner. **E**: There were fewer Ly-6C^hi^ cells during the asymptomatic clinical course (10 dpi) in mice that will develop the disease than in those that will remain asymptomatic. **F**: ROC curve analysis showing that the highest discriminatory ability when measuring Ly-6C^hi^ cells was found when classifying mild as opposed to severe EAE mice using the median SI. **G**: Cut-off values for the Ly-6C^hi^ cells/myeloid cells to predict a future mild or severe EAE clinical courses. A Student’s *t* test was used to compare the groups of animals and the mean (± SEM) are represented, at *P < 0.05. The statistical analyses were carried out using the Spearman correlation coefficient (n= 16 mice).

We addressed the discriminating power of the percentage of Ly-6C^hi^ cells at onset to assess the risk of developing mild or severe EAE at the peak of the disease according to the different parameters analyzed. Ly-6C^hi^ cells/myeloid cells presented a modest discriminating power to assess the risk of being mild/severe EAE mice based on the median value of the maximum clinical score at peak (AUC–area under the curve– 0.800, 95% CI –confidence interval– 0.514-1.026; p = 0.061) and the median value of the accumulated clinical score (AUC 0.813, 95% CI 0.571-1.054; p <0.05: Fig. 6F). By contrast, the level of Ly-6C^hi^ cells has a high discriminatory power when classifying mild or severe EAE mice using the median of the SI (AUC 0.964, 95% CI 0.875-1.052; p <0.01: Fig. 6F). In fact, we established cut-off values for the percentages of Ly-6C^hi^ cells/myeloid cells showing that values higher than 6.83% (100% specificity, 95% CI 59.04%-100.0%; 77.78% sensitivity, 95% CI 39.99%-97.19%) and lower than 5.47% (100% specificity, 95% CI 66.37%-100.0%; 85.71% sensitivity, 95% CI 42.13%-99.64%) at onset will be indicative of a future mild or severe EAE courses, respectively (Fig. 6G). In summary, these results suggested that the Ly-6C^hi^ cell content in the blood at the onset of the disease is a good predictor of the future EAE clinical course.

To check whether measuring of Ly-6C^hi^ cells in the peripheral blood would also be useful to reveal the future recovery of the same cohort of EAE mice, we evaluated the abundance of these cells at the peak of the disease and we related it to clinical parameters associated with the resolution of the clinical signs. We observed that the higher level of peripheral blood Ly-6C^hi^ cells at the peak of clinical course, the faster and greater recovery of EAE mice (Fig. 7A-C). In parallel, the peripheral Ly-6C^hi^ cells load at the disease peak was associated to a milder accumulated clinical score during the chronic phase (Fig. 7D). Moreover, the level of Ly-6C^hi^ cells/myeloid cells did not discriminate the risk of developing mild/severe course in EAE mice at the end of the recovery phase based on the median value of the recovery index (AUC 0.730, 95% CI 0.435-0.970; p = 0.172) and the median value (67 %) of the clinical score recovery (AUC 0.781, 95% CI 0.542-1.020: p = 0.058). However, the abundance of Ly-6C^hi^ cells at the peak of EAE showed a high discriminatory power using 50% as the recovery threshold (AUC 0.948, 95% CI 0.836-1.061; p <0.05: Fig. 7E). We established a cut-off value of 5.24% Ly-6C^hi^ cells/myeloid cells to be indicative of a future mild or severe EAE recovery course, respectively. As such, values higher than the cut-off showed 100% specificity (95% CI 29.24%-100.0%) and 92.31% sensitivity (95% CI 63.97%-99.81%), whereas values below the same cut-off showed a 92.31% specificity (95% CI 63.97%-99.81%) and 100% sensitivity (95% CI 29.24%-100.0%: Fig. 7F). These data reinforced the potential value of Ly-6C^hi^ cells as predictors not only of clinical course severity in the pro-inflammatory phase but also, of the partial recovery from clinical signs after the peak of the EAE course.

**Figure 7:**
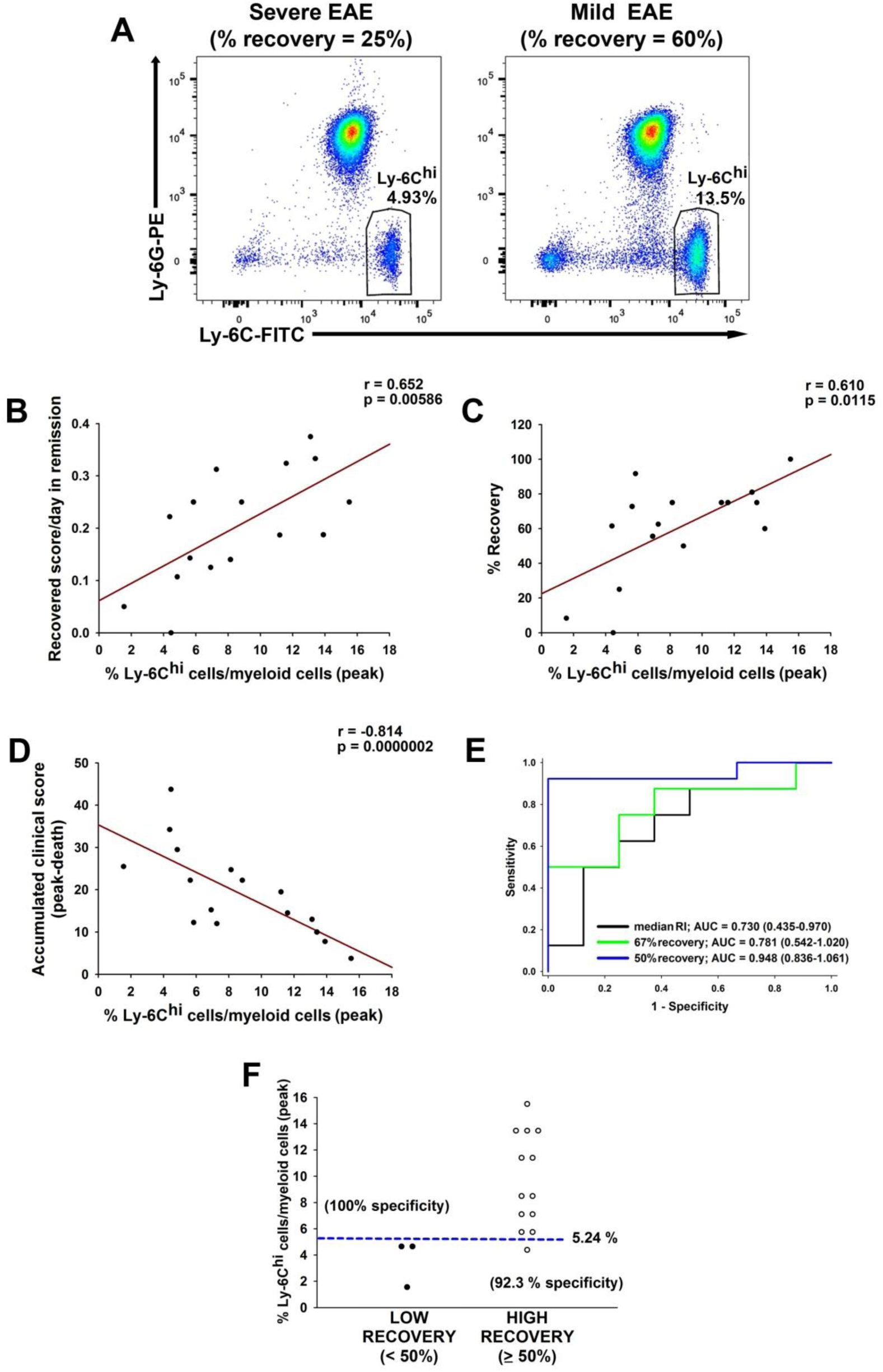
The higher abundance of Ly-6C^hi^ cells at the peak of the EAE clinical course is indicative of greater symptomatic recovery. **A**: Representative flow cytometry plots showing the percentage of Ly-6C^hi^ myeloid cells at the peak of the symptoms in EAE mice with a severe (left panel) or mild (right panel) clinical course. **B-D**: The abundance of Ly-6C^hi^ cells in the peripheral blood at the peak of the disease was directly correlated with recovery from the peak to the end of the follow-up (**B**), the percentage of symptomatic recovery (**C**) and the accumulated clinical score at the end of the clinical course (**D**). **E**: ROC curve analysis showed that the highest discriminatory power for Ly-6C^hi^ cells to classify the clinical course of EAE as mild or severe was observed using 50% recovery as the threshold. **F**: Cut-off value of Ly-6C^hi^ /myeloid cells at the peak of the disease to predict the future high recovery during the EAE clinical course. A Spearman test was carried out for the correlation analysis (n = 16 mice).

### The blood Ly-6C^hi^ cell content at the onset of the EAE clinical course is indicative of less CNS damage at the peak of the disease

Both demyelination and axonal damage are histopathological features related to the spinal cord abnormalities during the EAE clinical course and both are inversely correlated to the splenic M-MDSC content at peak (*16*). We sought to determine if there was a correlation between Ly-6C^hi^ cells in the peripheral blood at EAE onset and spinal cord pathology at the peak of disease. To this end, a second cohort of 10 EAE animals was used in a correlation analysis, confirming the correlation between Ly-6C^hi^ cells in the peripheral blood at the onset of the disease and the SI (r = -0.726; p < 0.05), the score at the peak of the disease (r = -0.778; p < 0.01) and the accumulated clinical score in this cohort (r = -0.65; p < 0.05). There was less destruction of myelin in the spinal cord of mice with mild clinical course than in those with severe EAE (Fig. 8A-B). Interestingly, higher abundance of blood Ly-6C^hi^ cells was linked to smaller areas of damaged myelin, as well as with the percentage it represented within the whole white matter (Fig. 8C-D). In parallel, axonal damage was less prominent in mild EAE mice than in those with a more severe clinical course (Fig. 8E-F), and the higher abundance of Ly-6C^hi^ cells in the peripheral blood of EAE mice at disease onset, the lower degree of axonal damage within the white matter (Fig. 8G) or within the infiltrated area (Fig. 8H).

**Figure 8:**
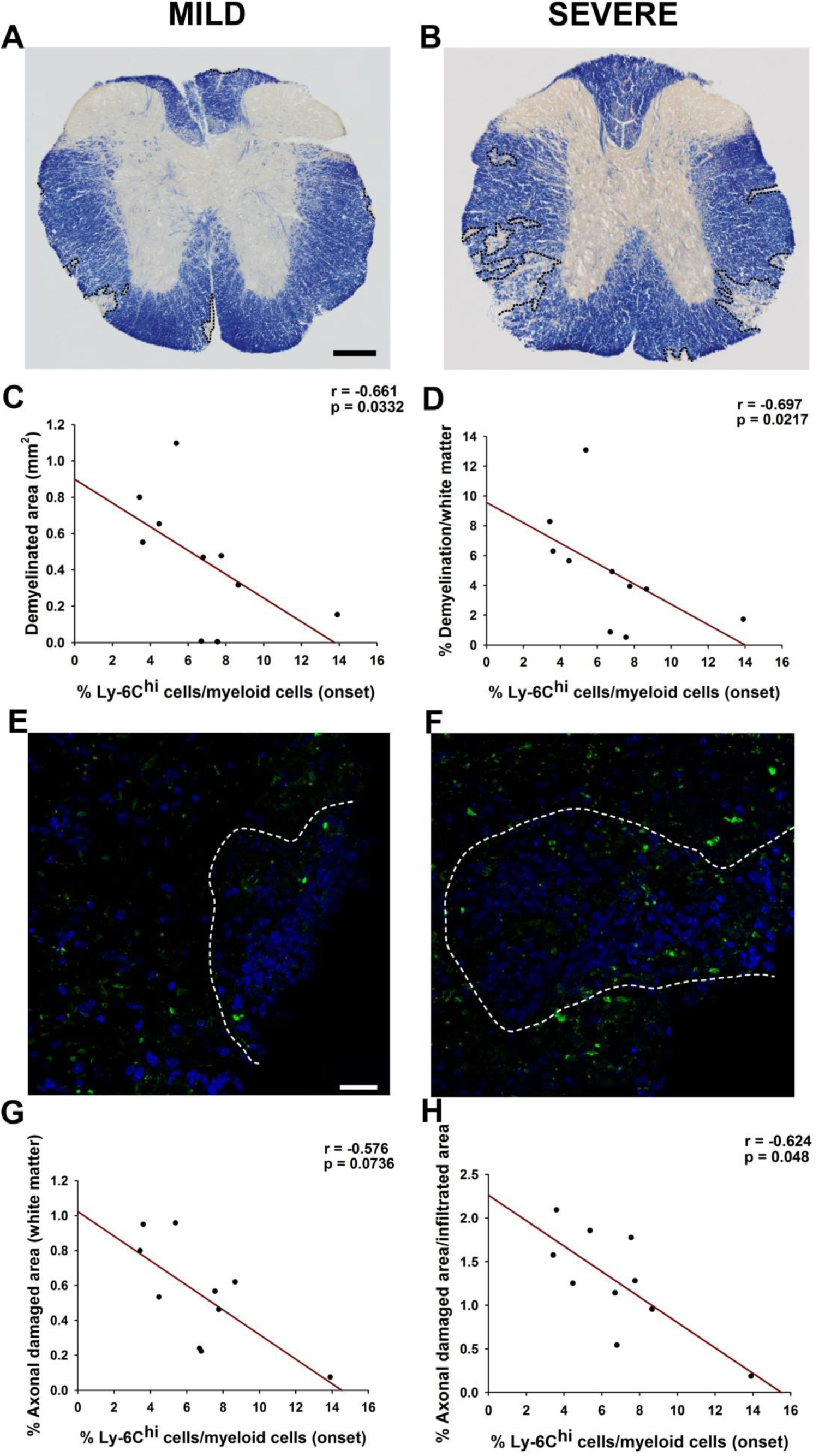
The Ly-6C^hi^ cell content in the peripheral blood at the onset of symptoms is indicative of lower CNS damage. **A-D**: Representative panoramic views of myelin staining with eriochrome cyanine of the spinal cord from a mild (**A**) or a severe (**B**) EAE mouse. The abundance of Ly-6C^hi^ cells at the onset of the clinical course was inversely correlated with the demyelinated area (**C**, **D**). **E-H**: SMI-32 staining showed more axonal damage in the spinal cord from severe (**F**) than in mild (**E**) EAE mice. Ly- 6C^hi^ cell abundance at the onset of the disease was inversely correlated with the extent of axonal damage (**G**, **H**). The statistical analyses were carried out using a Spearman correlation coefficient (n = 10 mice). Scale bar: A-B = 100 µm; E-F = 25 µm

In conclusion, the presence of a higher Ly-6C^hi^ cell content in blood on the first day of clinical signs pointed to a lower degree of spinal cord tissue affectation at the peak of the disease, indicating that measuring these cells in peripheral blood could be a good predictor for the evolution of the EAE clinical score.

### M-MDSCs in MS patients are indicative of a better relapse recovery

In a final step, we were interested in the predictive capacity of M-MDSCs in the blood of MS patients at an early time point in their clinical course. MDSCs were classified as M-MDSCs in the peripheral blood mononueclear cells (PBMCs) of healthy controls (HCs) and MS patients as CD33^+^ HLA-DR^-/low^ CD14^+^ CD15^-^ cells (Fig. 9A) (*23*). To draw a parallel with observations in EAE models, MS patients with a first clinical episode suggestive of MS in the last year were enrolled for this study (see Table S2 for demographic data). Overall, M-MDSCs seemed to be more abundant in MS patients than HCs (Fig. 9B). Regarding MS patients, the abundance of M-MDSCs was independent of their age (r = -0.168, p = 0.304) or EDSS at baseline (r = 0.00265, p = 0.987). Interestingly, the higher M-MDSC abundance was related to shorter periods of time from relapse to sampling (Fig. 9C). Since symptoms occurring within a month after previous clinical manifestations of an MS attack were considered to be part of the same relapse (*30*), we split the MS patients into two groups: MS patients whose blood was collected ≤ 30 days (patients in relapse) or more than 30 days after relapse (patients in remission; Table S2). MS patients during relapse but not MS patients in remission had a higher percentage of M-MDSCs than HCs (Fig. 9D), indicating that the proportion of M-MDSCs was higher close to the inflammatory episode.

**Figure 9:**
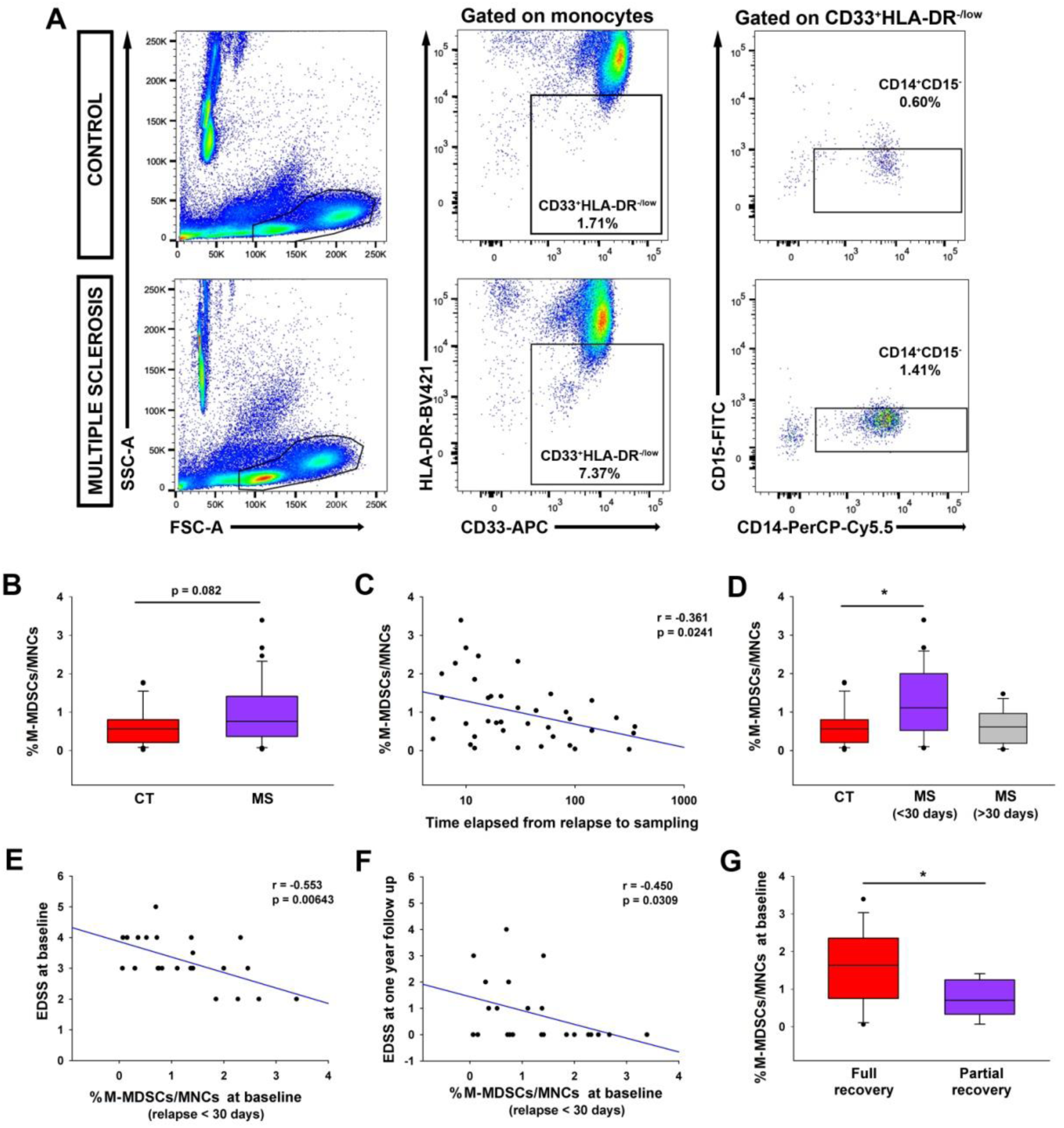
The abundance of M-MDSCs in the peripheral blood of MS patients is indicative of a better relapse recovery in MS. **A:** Representative flow cytometry plots for M-MDSCs in human PBMC samples. In both control and MS patients, live MNCs were gated after removing cellular aggregates and death cells based in Zombi-NIR expression. Next, immature myeloid cells were gated from monocyte subpopulation as CD33^+^ HLA-DR^−/low^ populations. CD33^+^ HLA-DR^−/low^ cells were assessed for CD14 and CD15 expression to identify the M-MDSC subset defined as CD14^+^ CD15^-^ (in this panel, the percentage of M-MDSCs refers to MNCs). **B**: The abundance of M-MDSCs measured in MS patients at an early stage of their disease course seemed to be higher than controls, although no significant differences were seen. **C:** The M-MDSC load in human peripheral blood was inversely correlated with the time elapsed from the first relapse to sampling. **D**: M-MDSCs were more abundant in MS patients at the time of their first referred relapse (MS-R, <30 days after the relapse) than in controls. **E-F:** In the sub-cohort of MS patients in relapse who were followed up for one year, the abundance of M-MDSCs at baseline was inversely correlated with the EDSS at baseline (**E**) and with the EDSS one year later (**F**). **G**: MS patients at relapse with full recovery (with an EDSS of 0 at 12 months) had a higher proportion of M-MDSCs than those with partial recovery. For correlation analysis a Spearman test was carried out (Control in B n= 26; MS in B and C n= 39; MS ≤30 days in D-G n = 23; MS >30 days in D n = 16; Total recovery n = 14 and partial recovery n = 9 MS patients in G). Comparative analysis in D was carried out with ANOVA and Student’s *t* test was used to compare groups of patients in B and G. The mean (± SEM) are represented, at *p < 0.05.

To establish the relationship between M-MDSCs and disease course, MS patients in relapse were followed up during one year. In this sub-cohort (Table S3), the M-MDSC content at baseline was still independent of the age (r = -0.214, p = 0.324). Importantly, M-MDSC abundance was associated to a less disability, i.e. the higher level of M-MDSCs at baseline the lower EDSS at sampling (Fig. 9E) and at one year follow-up (Fig. 9F). Moreover, there was a significant direct correlation with the percentage of EDSS recovery between both time points of sampling (r = 0.426; p < 0.05). Indeed, M-MDSCs were enriched in MS patients with a full relapse recovery (an EDSS of 0 at 12 months) than those who experienced partial recovery (p < 0. 05: Fig. 9G). Conversely, the proportion of M-MDSCs in the blood at one year follow-up was independent of the EDSS at that time point (r = -0.127, p = 0.561), nor was there any difference in the proportion of M-MDSCs between fully or partially recovered patients (Full recovery: 2.695 ± 0.613; Partial recovery: 2.247 ± 0.743; p = 0.395).

In summary, the data obtained clearly indicate that M-MDSCs were enriched in MS patients close to the relapse, and the high M-MDSC content is associated to a less disability at the time of sampling and to a better recovery after 12 months of follow-up.

## DISCUSSION

The data presented here show for the first time the presence of infiltrated myeloid cells with M-MDSC phenotype in human tissue from MS patients, mainly in areas with a high inflammatory activity. Indeed, we detected a direct correlation between the lower abundance of M-MDSCs in these areas and both the shorter age at death and disease duration in PPMS patients. Furthermore, we provided novel insights into the role of Ly-6C^hi^ cells/M-MDSCs as putative tools to gain information about the future clinical course of the EAE model in mice and of the evolution of EDSS in MS subjects at relapse. Our results show that when there is a higher abundance of Ly-6C^hi^ cells at the onset of EAE, the disease course will be milder. Interestingly, we corroborated that the abundance of M-MDSCs in blood samples from untreated MS patients at their first relapse is inversely correlated to the EDSS at baseline and after a one year follow-up.

Despite the enormous progress in developing DMTs for MS (*31*), a better understanding of the mechanisms driving the heterogeneity in the clinical course of the disease is crucial for the future prediction of disease progression, and the prompt and precise improvement of the disease by early and accurate treatments. The suppressive function of immunoregulatory cells such as Treg is closely related to pathological progression (*32, 33*). Furthermore, previous studies into MS using human brain samples demonstrated that the neuronal pathological changes and inflammation are closely related, not only in SPMS but also in PPMS patients, as they both display similar inflammatory activity (*7, 34, 35*). These observations highlight the role of the immune response in a subset of progressive MS patients, as confirmed by the approval of the immunomodulatory treatment ocrelizumab as the first EMA and FDA approved drugs for the treatment of PPMS (*36*). M-MDSCs are less abundant in SPMS patients than in RRMS patients and fail to suppress T-cell proliferation (*22*). Thus, the lack of correlation between the presence of M-MDSCs and the duration of the clinical courses of SPMS might be due to weaker immunosuppression or to exhaustion from repeated and more inflammatory attacks. Alternatively, the final distribution and probably the activity state of M-MDSCs in the CNS of SPMS may be the result of their modification after different therapeutic treatments during the relapsing-remitting phase. Indeed, this has been extensively described for cells of the innate immune response in the blood of MS patients (*37*– *41*), including MDSCs (*23, 42*). For the first time we show a correlation between the presence of regulatory myeloid cells (i.e. M-MDSCs) and the disease duration of PPMS. In our studies, the lower the abundance of M-MDSCs in areas with a high inflammatory activity, the shorter the disease course of PPMS patients. Furthermore, the ratio between CD14^+^ HLA-DR^hi^ pro-inflammatory macrophages and M-MDSCs (IIR) was inversely correlated with the disease duration in PPMS, suggesting that the dampened inmunoregulatory context might worsen the disease progression in these patients. Since all PPMS samples in the UK MS and Dame Ingrid V Allen tissue banks were collected before therapies for PPMS were available (2019), the M-MDSC distribution in these patients should reflect the distribution of immune cells at the end of the unmodified natural history of the disease. This point is supported by the identification of a similar direct correlation between the density of M-MDSCs and the age of PPMS patients. In this sense, the low density of M-MDSCs in high inflammatory regions of PPMS patients with shorter disease durations reinforces the notion that abnormalities in regulatory mechanisms may affect the clinical course. Our results may open the door for the future use of M-MDSCs as biomarkers of more benign/severe PPMS courses, which would have important consequences for the design of new clinical trials in this particular patients’ group as well as treatment decision-making by patients.

The data presented here show an inverse correlation between the levels of myeloid Ly-6C^hi^ cells in the peripheral blood of mice at the onset of the clinical signs and the severity of the disease course. In order to corroborate the parallels in Ly-6C^hi^ cell content, the severity of the disease course and the CNS damage, we analyzed the spinal cord of mice with EAE. A higher abundance of Ly-6C^hi^ cells at disease onset was inversely correlated with CNS damage, which included less destruction of myelin and axonal damage. Murine M-MDSCs from immune organs and infiltrated spinal cord of EAE mice share the same immunophenotype than the so-called Ly-6C^hi^ cells (*12, 28*). Interestingly, they may play different roles depending on the phase of the disease when they are isolated: Ly-6C^hi^ cells have pro-inflammatory activity at the onset of the disease whereas they behave as M-MDSCs when they are isolated at the peak of the clinical course (*28, 43, 44*). To our knowledge, circulating M-MDSCs are indistinguishable from inflammatory Ly-6C^hi^ monocytes in the peripheral blood (*14, 19*). For all the abovementioned data, circulating Ly-6C^hi^ cells appears as the only appropriate cell subset to be analyzed in order to validate M-MDSCs as a predictive tool for EAE severity. It has been described that almost 98% of the Arg-I^+^ anti-inflammatory cells that infiltrate the CNS at the peak of the clinical course were CCR2^+^-invading cells (*45*). Futhermore, 50% of infiltrating Arg-I^+^ macrophages in the CNS at the peak derived from pro-inflammatory iNOS-expressing cells invading this area at the onset of the clinical course (*46*). Taking together, it could be thought that the higher Ly-6C^hi^ peripheral blood cell content at the onset of the disease, the higher density of anti-inflammatory cells found at the peak of the symptoms. Previous studies illustrated how the anti-inflammatory environment promoted by the M2 polarization of microglia/macrophages may help ameliorate EAE progression and promote remyelination (*47, 48*). Furthermore, the capacity of Ly-6C^hi^ cells to change their activity was reflected by the shift from pro-inflammatory to a clear anti-inflammatory activity following pharmacological treatment with IFN-β (*18*), suggesting that the presence of this malleable cell type at the onset of EAE will result in a higher abundance of M-MDSCs with immunosuppressive activity at the peak of the disease. It is beyond the scope of this work to determine whether these Ly-6C^hi^ cells found at disease onset are the same cells as those detected at the disease peak. However, the crucial role of pro-inflammatory Ly-6C^hi^ cells in the effective repair at later stages of chronic inflammatory pathologies was reported recently (*49*). In parallel with these results, the stronger presence of Ly-6C^hi^ cells at the onset of the clinical signs in the EAE model might help enhance the repair mechanisms involved in the later stages of the clinical course of EAE. In support of this, it was recently described that M-MDSCs promote remyelination in EAE by enhancing oligodendrocyte precursor cell (OPC) survival, proliferation and differentiation (*50*), suggesting the crucial role of M-MDSCs not only in resolving inflammation but also in promoting tissue regeneration and effective recovery.

Beyond searching for biomarkers that help us to predict the severity of the MS clinical course, a great deal of work has focused on tissue repair, since promoting remyelination has been directly associated with improved clinical recovery in animal models (*51–53*). However, a challenge for remyelination trials is the lack of a robust biomarker of successful regeneration, since the specificity of high resolution imaging-based markers for myelin regeneration remains questionable (*54*). Recent advances in developing specific imaging tools to measure myelination *in vivo* have shed light on the potential effects of new pro-remyelinating therapies but also, on classifying patients depending on their individual remyelination potential (*55*). Similarly, the search for useful biological parameters to predict future symptomatic recovery needs to be addressed. In this sense, our observations show that the number of Ly-6C^hi^ cells at the peak of the symptoms is also correlated with recovery in mice with EAE. Indeed, more than 5.24% Ly-6C^hi^ cells within the myeloid cell subset at the time of the maximum clinical score seems to represent a threshold that significantly discriminates those mice that will recover at least 50% of the clinical impairment. To our knowledge, for the first time our data indicate that cells with a M-MDSC phenotype (Ly-6C^hi^ cells) can be used as putative biological tools to gain information about the recovery from symptoms, which is reinforced by our data regarding the inverse relationship between M-MDSCs in the blood of MS patients at sampling and their EDSS one year later.

The immunotherapies currently available are largely insufficient to prevent the accumulation of chronic disability in the progressive phases of MS and hence, researchers are currently focusing their investigations on alternative therapeutic options, such as restoring myelin to protect against neurodegeneration (*51, 52, 56*). Given the accumulating evidence of the beneficial role of MDSCs, not only as powerful suppressors of T cell activity but also as important regulatory agents for recovery from immunological insults by promoting Tregs (*57–60*), it seems logical to consider these cells as contributors to the protection of the key cellular components of the CNS in the context of MS (i.e.: myelin/oligodendrocytes and axons/neurons). Hence, a stronger presence of M-MDSCs at the peak of the disease seems to be related to a better resolution of inflammation and tissue repair. Given that Treg cells are required for efficient OPC differentiation during remyelination (*61*), the abundance of M-MDSCs may help increase the Treg population, which in turn would support myelin regeneration. It is important to note that we recently showed direct effects of M-MDSCs on survival and proliferation of OPCs, also showing them to be potent promoters of OPC differentiation towards mature phenotypes (*50*). Hence, this regulatory cell population should be considered for the development of future therapeutic tools focused on CNS repair. In parallel, this study shows that the density of M-MDSCs is significantly higher in ALs and the rAIL, regions in which it has been shown that remyelination can potentially occur (*27, 62, 63*). The protective role of the innate immune system has been described previously and is crucial for promoting myelin repair after CNS damage. Indeed, the resolution of pro-inflammatory microglia activation through a switch to a pro-regenerative state initiates remyelination (*47, 64, 65*). Thus, we propose that the presence of M-MDSCs in areas with the possibility to be repaired may help enhance the remyelination of these regions, not only by resolving the immune response by T cell immunosuppression but also, by indirectly enhancing repair mechanisms, such as those mediated by Treg cells.

Finally, we found that the abundance of M-MDSCs in blood samples from untreated MS patients at relapse is correlated with a lower EDSS at baseline and after a one year follow-up. Hence, the analysis of this regulatory cell population suggests a role of M-MDSCs as a future biomarker of MS clinical progression, though the stratification based on DMTs needs for further analysis in larger cohorts. It is important to note that MDSCs were recently shown to have certain relevance in terms of the recovery of the MS clinical course, whereby the expansion of MDSCs in glucocorticoid-treated MS patients may help alleviate the acute phase of the disease (*42*). However, these promising results about the role of MDSCs as biomarkers of MS severity should be followed-up in a long-standing and larger cohort of MS patients.

In summary, the results we obtained from studying human samples together with observations in the EAE model suggest that M-MDSCs may be a reliable measure predicting the clinical course of relapses and disease evolution, opening the door for their consideration as a future bioindicator for relapse management. As such, the therapeutic use of M-MDSCs could be considered as a strategy for improving immunoregulatory mechanisms as well as myelin repair to alleviate the disease course.

## MATERIALS AND METHODS

### Human tissue

*Post-mortem* cortical snap-frozen brain tissue blocks were analyzed from MS patients of both sexes and with no history of neuropsychiatric disease, as well as that from 6 matched human controls (Ct: Table 1). The study included 33 cases of MS comprising secondary progressive MS (SPMS, n = 20) and PPMS (n = 13). All the cortical blocks contained both gray matter and white matter. The UK MS Tissue Bank randomly provided us with at least three different blocks from the cerebral cortex with white matter lesions for each MS patient (labeled as MS), and two cortical blocks from each selected control. Formalin-fixed paraffin-embedded (FFPE) MS tissue blocks (labeled as MSD) were also provided by Prof. Denise Fitzgerald from the Dame Ingrid V. Allen tissue collection (Belfast, UK). No demyelinated lesions were detected within the gray matter in any of the blocks analyzed. The brain tissue was cut into cryosections (MS: 10 µm, Leica cryostat; MSD: 6 µm Leica, RM2255 microtome) for immunohistochemical analysis.

Classification of MS patients based on the clinical course was performed according to the median of the disease duration of SPMS and PPMS (SPMS: 16 years, inter quartile range-IQR 13,5-34,5; PPMS: 17 years, IQR 11-28,5; Table 1). Short clinical course was considered when disease duration was ≤ 16 years in SPMS (14 years, IQR 12-15) or ≤ 17 years in PPMS (13 years, IQR 10-15). On the other hand, SPMS patients with a long clinical course were considered if disease duration was > 16 years (35 years, IQR 33-41) while in PPMS patients the long clinical course was determined when it was > 17 years (30 years, IQR 27-35).

### Immunohistochemistry and eriochrome cyanine for myelin staining

The cryostat sections were fixed in 4% paraformaldehyde (PFA: Sigma-Aldrich) in 0.1 M Phosphate Buffer, pH 7.4 (PB) for 1 h at room temperature (RT). After several rinses with PB, the sections were incubated for 1 h at RT in incubation buffer: 1X Phosphate buffer saline (PBS) containing 5% normal donkey serum and 0.2% Triton X-100 (Merck). Subsequently, immunohistochemistry (IHC) or immunofluorescence (IF) staining was performed by incubating sections overnight at 4°C with the following primary antibodies: rabbit anti-CD11b (1:100; Abcam, ab133357); rabbit anti-TMEM119 (1:50; Sigma-Aldrich, HPA0518870); biotinylated sheep anti-CD14 (1:25; R&D, BAF383), mouse anti-CD15 (1:25; Agilent, ISO62, carb-3 clone) and mouse anti-human leukocyte antigen (HLA)-DR (1:200 for IHC, 1:100 for IF; Agilent, M0746, TAL.1B5 clone). Appropriate fluorescent-tagged (1:1000, Invitrogen) or biotinylated (1:200; Vector Labs) secondary antibodies were used. Detection of TMEM119 was visualized using TSA signal amplification system (Tyramide SuperBoost™ kit, Invitrogen). The IHC reaction was developed using the Vectastain Elite ABC reagent (Vector Labs) and the peroxidase reaction product was visualized with 0.05% 3,3′-diaminobenzidine (Sigma-Aldrich) and 0.003% H_2_O_2_ in 0.1 M Tris-HCl, pH 7.6. The reaction was monitored under the microscope and terminated by rinsing the slides with PB. Fluorescent Hoechst 33342 staining (10 µg/ml: Sigma-Aldrich) was used to label the cell nuclei. Negative stain controls were included and in all conditions the absence of the appropriate antibodies yielded no signal.

To visualize myelin after HLA-DR immunostaining in tissue from snap-frozen brain blocks, eriochrome cyanine (EC) staining was carried out, as described previously (*13*). The sections were air-dried overnight at RT and for 2 h at 37 °C in a slide warmer. The sections were then placed in fresh acetone for 5 min and air-dried for 30 min, before they were stained in 0.5% EC for 1 h and differentiated in 5% iron alum and borax-ferricyanide for 10 and 5 min, respectively (briefly rinsing the sections in tap water between each step). After washing with abundant water, correct differentiation was assessed under the microscope whereby the myelinated areas were stained blue and the demyelinated areas appeared white-yellowish. The stained sections were dehydrated and mounted for preservation at RT.

Triple chromogenic immunohistochemistry on FFPE sections (from MSD tissue blocks) was performed with a triple stain IHC kit (DAB, AP/Red & HRP/Green) from Abcam (ab183286) which allowed red-green colocalisation using a modified protocol. FFPE sections were dewaxed in clearene and rehydrated through a graded series of alcohols. Heat-induced epitope retrieval was conducted in a steamer for 1 h whilst slides were incubated in sodium citrate buffer (pH 6.0). Endogenous peroxidases were blocked with 0.3% hydrogen peroxide in methanol before blocking with 5% normal goat serum in Tris buffered saline (TBS, pH 7.4). Slides were then incubated with mouse anti-CD15 primary antibody (1:50; Agilent, ISO62, carb-3 clone) in blocking solution overnight. Slides were washed in TBS before EnVision anti-mouse HRP secondary antibody (Agilent) was applied for 30 min at RT. Antibody binding was visualised with 3, 3’-diaminobenzidine (DAB; Immpact DAB; Vector) as chromogen. To prevent cross-reactivity, slides were heated to 80°C in supplied antibody blocker solution, and then incubated with Blocker A and B according to the manufacturer’s protocol. Slides were washed in TBS before incubation with mouse anti-HLA-DR antibody (1:200; Agilent, M0746, TAL.1B5 clone) and sheep anti-CD14 antibody (1:50; R&D, BAF383) in blocking solution overnight at 4°C. After washing, tissue sections were incubated with kit-supplied anti-mouse AP secondary antibody for 30 min at RT. Followed further washing, sections were incubated with an ABC peroxidase-linked reporter system, (Vector Laboratories) for 30 min at RT. Detection of HLA-DR was visualised with Permanent Red chromogen and counterstained with Gill’s haematoxylin No. 2 (GHS232; Sigma) diluted 1:10. Slides were briefly washed and detection of CD14 was visualised by applying Emerald Green chromogen for 5 min at RT. Slides were rapidly cleared and mounted with supplied mounting medium according to the manufacturer’s instructions. Please note that for quantitative analysis, all slides were stained in the same experimental run. Negative and single stain controls were included and in all instances the absence of the relevant antibodies yielded no signal.

### Classification of MS lesions

White matter MS lesions were classified according to demyelination and the distribution of the HLA-DR^+^ cells, as previously described (*8, 65–67*). Active lesions (AL) had abundant and evenly distributed HLA-DR^+^ cells within the demyelinating lesion, cells that were mostly ameboid. Mixed active/inactive lesions (AIL) are demyelinated regions characterized by a hypocellular lesion center (cAIL) and a rim enriched with HLA-DR^+^ cells at the lesion border (rAIL). Finally, inactive lesions (IL) contained very few HLA-DR^+^ cells within the complete demyelinating lesion. According to the MS classification, the AL and the rAIL were considered regions with a high inflammatory activity. Alternatively, the areas with weak inflammatory activity included the cAIL and IL.

### Cell counting in human tissue

The IHC staining for HLA-DR, CD14 and CD15 was used for the quantification of M-MDSCs in serial sections of each case and lesion. To measure HLA-DR fluorescence intensity, photomicrographs of the MS lesions were acquired as a mosaic of 20X magnification images captured on a confocal microscope equipped with a resonant scanning system (SP5: Leica), quantifying the fluorescence intensity of the cells using the ImageJ software. A low level of HLA-DR fluorescence was established by measuring HLA-DR staining in one hundred CD14^+^ CD15^-^ cells, among which there were 25 cells with no HLA-DR immunostaining, 25 cells with very faint staining (HLA-DR^low^) and 50 cells with mild-to-strong immunostaining (HLA-DR^int/high^: Fig. S1A). When the maximum fluorescence intensity was quantified in each HLA-DR cell population, a receiver operating characteristic (ROC) curve analysis was performed to find the optimal cut-off to accurately classify HLA-DR^low^ cells. Once this threshold was calculated, only those cells with HLA-DR fluorescence intensity under the cut-off value were considered to quantify the M-MDSC density (Fig. S1B). The density of M-MDSCs was obtained by manually counting of CD14^+^ HLA-DR^-/low^ CD15^-^ cells within the MS lesions, using 5-15 fields of the area of interest at a magnification of 20X, depending on the size of the lesions (SP5: Leica). In the same fields, the quantification of CD14^+^ HLA-DR^hi^ CD15^-^ cells was manually obtained to calculate the relative abundance of immunoregulatory cells in the MS lesion by using inflammatory-immunoregulatory ratio (IIR), i.e. the density of CD14^+^ HLA-DR^hi^ cells/density of M-MDSCs .

Colour deconvolution was performed to quantify M-MDSCs (HLA-DR^-/low^ CD14^+^ CD15^-^ cells) in the MS lesions from FFPE tissue from the Dame Ingrid V. Allen tissue collection (Belfast, UK). The colour deconvolution plugin for Fiji implements stain separation with Ruifrok and Johnston’s method previously described (*68*). The plugin allows us to transform each single staining from the multiple immunolabelling into a separate channel to analyze the pictures and quantified the density of MDSCs in different MS lesions as described above (Fig. S1A-B).

### Induction of EAE

Female six-week-old C57/BL6 mice were purchased from Janvier Labs. Chronic progressive EAE was induced by subcutaneous immunization with 200 µl of the Myelin Oligodendrocyte Glycoprotein (MOG_35-55_) peptide (200 µg: GenScript), emulsified in complete Freund’s adjuvant containing heat inactivated *Mycobacterium tuberculosis* (4 mg/ml: BD Biosciences). Immunized mice were intravenously administered Pertussis toxin (250 ng/mouse: Sigma-Aldrich) through the tail vein on the day of immunization and 48 h later. EAE was scored clinically on a daily basis in a double-blind manner: 0, no detectable signs of EAE; 0.5, half of the tail paralyzed; 1, completely paralyzed tail; 1.5, partial weakness or altered hind limb movement; 2, weakness or partial unilateral hind limb paralysis; 2.5, partial bilateral hind limb paralysis; 3, complete bilateral hind limb paralysis; 3.5, complete bilateral hind limb and partial paralysis of the forelimbs; 4, total paralysis of the forelimbs and hind limbs; 4.5, moribund; and 5, death.

Following ethical standards and regulations, humane end-point criteria were applied when an animal reached a clinical score ≥ 4, when a clinical score ≥ 3 was reached for more than 48 h, or whether signs of stress or pain were evident for more than 48 h, even if the EAE score was < 3. Stress was considered as the generation of sounds, stereotypic behavior, lordokyphosis, hair loss or weight loss superior to 2 g/day. All animal manipulations were approved by the institutional ethical committees (*Comité Ético de Experimentación Animal del Hospital Nacional de Parapléjicos*), and all the experiments were performed in compliance with the European guidelines for animal research (European Communities Council Directive 2010/63/EU, 90/219/EEC, Regulation No. 1946/2003), and with the Spanish National and Regional Guidelines for Animal Experimentation and the Use of Genetically Modified Organisms (RD 53/2013 and 178/2004, Ley 32/2007 and 9/2003, Decreto 320/2010).

The clinical parameters analyzed were defined as: i) the severity index (SI), quantified as the ratio between the maximal clinical score at peak and the disease duration [i.e.: days elapsed from the onset to the peak of the disease (*16*)]; ii) the accumulated clinical score was considered as the sum of the individual clinical scores from the day of onset or the peak of the disease, until the end of the clinical evaluation; iii) the percentage of recovery was determined as the following percentage: (the maximal clinical score at peak – the residual score in the plateau phase)*100/maximal clinical score at peak; and iv) the recovery index, as the absolute score recovered from the peak to the plateau phase/days elapsed from the peak to the end of the remission phase.

### Flow cytometry analysis of peripheral blood cells from EAE mice

A first cohort of 16 mice was used for flow cytometry analysis. Blood was collected from isoflurane-anesthetized mice with EAE at the onset of the disease (the first day when mice showed a clinical score ≥ 0.5) and at the peak of the clinical symptoms (the peak was defined as the first day a clinical score ≥ 1.5 was observed as a repeat score). Approximately 75 µL of blood was obtained from the orbital vein of each mouse and collected in 2% EDTA tubes. Erythrocyte lysis was then performed in a 15 ml tube with ACK lysis buffer: 8.29 g/L NH_4_Cl; 1 g/L KHCO_3_; 1 mM EDTA in distilled H_2_O at pH 7.4 (Panreac). Subsequently, the cells were resuspended in 25 µl of staining buffer: sterile PBS supplemented with 10% fetal bovine serum (FBS: Linus); 25 mM HEPES buffer (Gibco); 1 mM EDTA and 2% Penicillin/Streptomycin (P/S: Gibco). Fc receptors were blocked for 10 min at 4°C with anti-CD16/CD32 antibodies (10 µg/mL, BD Biosciences) and the cells were labeled for 30 min at 4°C in the dark with the corresponding antibody panel: anti-Ly-6C FITC (10 µg/mL, AL-21 clone, BD Biosciences), anti-Ly-6G PE (4µg/ml, 1A8 clone, BD Biosciences), anti-CD11b PerCP-Cy5.5 (4µg/ml, M1/70 clone, BD Biosciences), anti-MHC-II PE-Cy7 (4µg/ml, M5/114.15.2 clone, eBioscience), anti-CD11c APC (4µg/ml, N418 clone, eBioscience) and anti-F4/80 eFluor450 (4µg/ml, BM8 clone, eBioscience). The blood cells were then rinsed with staining buffer, recovered by centrifugation at 490g for 5 min at RT, fixed for 10 min at RT with 4% PFA, and after a centrifugation step, resuspended in PBS and finally analyzed in a FACS Canto II cytometer (BD Biosciences) at the Flow Cytometry Service of the *Hospital Nacional de Parapléjicos*. A total number of 30,000 events/mice were acquired in CD11b^+^ region. Data analysis was assessed using the FlowJo 10.6.2 software (FlowJo, LLC-BD Biosciences).

### Tissue extraction and histological analysis of EAE tissue

Ten mice with EAE from a second cohort of animals were used for histological analysis. Peripheral blood was collected from all the mice at the onset of the clinical signs and all the animals were sacrificed at the peak of the clinical course, when they were perfused transcardially with 4% PFA. The spinal cord of the mice was dissected out and post-fixed for 4 h at RT in the same fixative. After immersion in 30% (w/v) sucrose in PB for 12 h, coronal cryostat sections (20 μm thick, Leica) were thaw-mounted on Superfrost® Plus slides.

The same EC staining for myelin visualization was carried out as that used for the histopathology of human samples with the following modifications: the tissue was stained in 0.5% EC for 30 min, and differentiated in 5% iron alum and borax-ferricyanide for 10 and 5 min, respectively. Axonal damage was analyzed by immunohistochemistry to stain the non-phosphorylated form of the neurofilament protein (SMI-32). After several rinses with PB, the tissue was pre-treated for 15 min with 10% methanol in PB and the sections were pre-incubated for 1 h at RT in incubation buffer. Immunohistochemistry was performed by incubating the sections overnight at 4°C with the primary SMI-32 antibody (1:200, Covance), diluted in incubation buffer. After rinsing, the sections were then incubated for 1 h at RT with the corresponding fluorescent secondary antibody (1:1000, Invitrogen). The cell nuclei were then stained with Hoechst 33342 (10 µg/ml, Sigma-Aldrich), and the sections were mounted in Fluoromount-G (Southern Biotech). Negative stain controls were included and in all conditions the absence of the appropriate antibodies yielded no signal.

### Image acquisition and analysis of murine tissue

In all cases, 3 sections from each thoracic spinal cord (separated by 420 μm) were selected from 10 mice with EAE in the histological cohort. To measure demyelination, the EC stained spinal cord sections were analyzed on a stereological Olympus BX61 microscope, using a DP71 camera (Olympus) and VisionPharm software for anatomical mapping. Superimages were acquired at a magnification of 10X using the mosaic tool and analyzed with the Image J software, expressing the results as the percentage of white matter area with no signs of blue staining as well as the total demyelinated white matter area.

To quantify axonal damage, mosaic images from the whole spinal cord of each animal were obtained on a DMI6000B microscope (Leica). The area of axonal damage relative to the total area or the infiltrated area was analyzed with an *ad-hoc* plug-in designed by the Microscopy and Image Analysis Service at the *Hospital Nacional de Parapléjicos*. Briefly, after selecting the appropriate area (the infiltrated area relative to the whole section or to the whole white matter area), a threshold for immunofluorescence was established and SMI-32 immunostaining was assessed, presenting the result as an area (μm^2^).

### MS patient cohort for M-MDSC blood analysis

All MS patients in this study had MS according to the revised 2017 McDonald criteria, and the cohort included 35 untreated RRMS patients who had not received corticosteroids in the last 6 months and who experienced their first relapse up to one year before blood sampling (demographics of the cohort are shown in Table S2). A clinical relapse was defined as any new neurological dysfunction or worsening of existing symptoms, lasting more than 24 h in the absence of fever or infection. Disability was assessed by trained neurologists using Kurtzke’s Expanded Disability Status Scale (EDSS), both at the time of blood sampling and one year later. All MS patients were recruited at the Department of Neurology at *Hospital Universitario Virgen de la Salud* (Toledo) or at *Hospital General Universitario Gregorio Marañón* (Madrid, Spain), during routine clinical visits, or at presentation during visits to the Casualties Medical Service or Inpatient visit at each hospital. Peripheral blood samples were obtained from healthy volunteers recruited in the Research Unit of the *Hospital Nacional de Parapléjicos*. The study was approved by the *Comité Ético de Investigación Clínica con Medicamentos* (number 349) of the *Complejo Hospitalario de Toledo* and informed, written consent was obtained from all participants in accordance with the Helsinki declaration.

### Preparation of human peripheral blood mononuclear cells and flow cytometry analysis of M-MDSCs

Human peripheral blood mononuclear cells (PBMCs) were isolated by Ficoll density gradient centrifugation (GE-171440-02, Merck). Separated cells were subsequently collected from the interphase, washed with isolation buffer (2.23 g/L D-glucose, 2.2 g/L sodium citrate, 0.8 g/L citric acid, 0.5% BSA in PBS) and further centrifuged at 500g for 10 min at RT. The cell pellet was resuspended in FBS, counted and aliquoted 1:1 in FBS with 20% dimethyl sulfoxide (DMSO, Sigma-Aldrich). The samples were then stored in liquid nitrogen at -160 °C until use.

Freshly thawed PBMCs (1x10^6^) were washed with RPMI and stained with Zombie NIR Dye (Biolegend) following the manufacturer’s instructions. After 15 min at RT in the dark, the cells were washed with PBS and recovered at 490g for 5 min. Then, PBMCs were resuspended in 25 μl of staining buffer and incubated for 10 min at 4°C with Beriglobin (50 μg/ml: CSL Behring) to block the Fc receptors. Next, 25 µl of the antibody mixture diluted in staining buffer were added to the cells for an additional 30 min at 4°C in the dark. The antibody panel was made up of anti-CD15-FITC (1.25µl/test, HI98 clone), anti-CD14-PerCP-Cy^TM^5.5 (0.5µl/test, Mφ29 clone), anti-CD11b-PE-Cy7 (0.5µl/test, ICRF44 clone), anti-CD33 APC (1.25µl/test, WM53 clone) and anti-HLA-DR BV421 (0.5µl/test, G46-6 clone, all from BD Biosciences). PBMCs were washed with staining buffer at 490g for 5 min at RT, resuspended in 0.1% PFA diluted in PBS and stored at 4°C in the dark until the next day. The samples were analysed in a FACS Canto II cytometer (BD Biosciences) with the corresponding controls and 100,000 events were recorded in the mononuclear cells (MNCs) gate. Compensation controls were obtained by incubating each single antibody with the Anti-Mouse Ig, k/Negative Control Compensation Particle Set (CompBeads, BD Biosciences). Fluorescence Minus One controls (FMO) were set-up in pooled samples [including healthy controls (HCs) and MS patients] to avoid any variation between the experimental conditions. The data were analysed with FlowJo v.10.6.2 software (FlowJo, LLC-BD Biosciences).

### Statistical analysis

The data are expressed as the mean ± SEM and they were analyzed with SigmaPlot version 11.0 (Systat Software). To compare between all the different MS lesions, a one way ANOVA test was performed or its corresponding ANOVA on ranks, followed by the Tukey or Dunn *post-hoc* tests, respectively. The Student’s *t* test was used to perform two-by-two comparisons (Mann-Whitney *U* test for non-parametric data). Shapiro–Wilk normality tests were performed on all human MS tissue samples. Pearson or Spearman tests were used for correlations as appropriate. The ROC curve was quantified using the area under the curve (AUC) to evaluate the predictive value of MDSC abundance in categorizing disease severity of EAE mice (severe or mild). The minimal statistical significance was set at p < 0.05 and the results of the analysis were represented as: *, # p< 0.05; **, ## p<0.01; ***, ### p < 0.001.

## Acknowledgements

### Funding

This work was supported by the Instituto de Salud Carlos III-Spanish Ministerio de Ciencia e Innovación (PI18/00357 and RD16-0015/0019, both partially co-financed by F.E.D.E.R., European Union, “*Una manera de hacer Europa*”, and PI21/00302, co-financed by the European Union), Fundación Merck Salud, ARSEP Foundation, Esclerosis Múltiple España (REEM-EME-S5 and REEM-EME_2018), ADEMTO, ATORDEM and AELEM (to DC) and the Wellcome Trust (to DCF). DC, RL-G, MCO, IM-D, IPM, VG, RG-M, and VVdS are hired by SESCAM. JG-A was hired by RD16-0015/0019. CC-T hold a predoctoral fellow by the Instituto de Salud Carlos-III (FI19/00132, partially funded by Fondo Social Europeo “*El Fondo Social Europeo invierte en tu futuro*”). The authors would like to thank: Dr José Ángel Rodríguez-Alfaro and Dr Javier Mazarío, from the Microscopy Service of the *Hospital Nacional de Parapléjicos* for their assistance with the confocal imaging, and histological quantifications; and Teresa Hernández-Iglesias for her advice in the search and adaptation of immunological markers for histopathology of the human CNS.

### Authors’ contributions

MCO and RLG contributed equally to this work. MCO and RL-G performed most of the experiments. MCO was a major contributor in writing the manuscript. IM-D contributed to the analysis of blood samples. JG-A contributed with M-MDSC quantification in human samples. MN and CC-T performed part of the histopathological analysis in humans and mice, respectively. VVdS contributed to the design of the strategy and analysis of M-MDSC in blood samples. IP-M, RG-M, VG, HG, JMG-D and MLM-G carried out MS patient selection and clinical evaluation at baseline and during the follow-up. MN designed and carried out studies from the Dame Ingrid V. Allen tissue collection. DF oversaw work with tissue from the Dame Ingrid V Allen tissue collection and contributed to editing of the manuscript. DC designed the experiments, contributed to data analysis and was a major contributor in writing the manuscript. All authors read and approved the final manuscript.

### Data and materials availability

All data are available in the main text or the supplementary materials

## References

1. A. J. Thompson, S. E. Baranzini, J. Geurts, B. Hemmer, O. Ciccarelli, Multiple sclerosis. Lancet. 391 (2018), pp. 1622–1636.

2. R. M. Ransohoff, D. A. Hafler, C. F. Lucchinetti, Multiple sclerosis - A quiet revolution. Nat. Rev. Neurol. 11 (2015), pp. 134–142.

3. D. S. Reich, C. F. Lucchinetti, P. A. Calabresi, Multiple Sclerosis. N. Engl. J. Med. 378, 169–180 (2018).

4. C. A. Dendrou, L. Fugger, M. A. Friese, Immunopathology of multiple sclerosis. Nat. Rev. Immunol. 15 (2015), pp. 545–558.

5. M. Tintore, A. Vidal-Jordana, J. Sastre-Garriga, Treatment of multiple sclerosis — success from bench to bedside. Nat. Rev. Neurol. 15 (2019), pp. 53–58.

6. M. Filippi, M. A. Rocca, O. Ciccarelli, N. De Stefano, N. Evangelou, L. Kappos, A. Rovira, J. Sastre-Garriga, M. Tintorè, J. L. Frederiksen, C. Gasperini, J. Palace, D. S. Reich, B. Banwell, X. Montalban, F. Barkhof, MRI criteria for the diagnosis of multiple sclerosis: MAGNIMS consensus guidelines. Lancet Neurol. 15 (2016), pp. 292–303.

7. R. Magliozzi, O. W. Howell, R. Nicholas, C. Cruciani, M. Castellaro, C. Romualdi, S. Rossi, M. Pitteri, M. D. Benedetti, A. Gajofatto, F. B. Pizzini, S. Montemezzi, S. Rasia, R. Capra, A. Bertoldo, F. Facchiano, S. Monaco, R. Reynolds, M. Calabrese, Inflammatory intrathecal profiles and cortical damage in multiple sclerosis. Ann. Neurol. 83, 739–755 (2018).

8. S. Luchetti, N. L. Fransen, C. G. van Eden, V. Ramaglia, M. Mason, I. Huitinga, Progressive multiple sclerosis patients show substantial lesion activity that correlates with clinical disease severity and sex: a retrospective autopsy cohort analysis. Acta Neuropathol. 135, 511–528 (2018).

9. K. Kirschbaum, J. K. Sonner, M. W. Zeller, K. Deumelandt, J. Bode, R. Sharma, T. Krüwel, M. Fischer, A. Hoffmann, M. C. Da Silva, M. U. Muckenthaler, W. Wick, B. Tews, J. W. Chen, S. Heiland, M. Bendszus, M. Platten, M. O. Breckwoldt, In vivo nanoparticle imaging of innate immune cells can serve as a marker of disease severity in a model of multiple sclerosis. Proc. Natl. Acad. Sci. U. S. A. 113, 13227–13232 (2016).

10. M. C. Gjelstrup, M. Stilund, T. Petersen, H. J. Møller, E. L. Petersen, T. Christensen, Subsets of activated monocytes and markers of inflammation in incipient and progressed multiple sclerosis. Immunol. Cell Biol. 96, 160–174 (2018).

11. F. Veglia, M. Perego, D. Gabrilovich, Myeloid-derived suppressor cells coming of age review-article. Nat. Immunol. 19 (2018), pp. 108–119.

12. B. Zhu, Y. Bando, S. Xiao, K. Yang, A. C. Anderson, V. K. Kuchroo, S. J. Khoury, CD11b + Ly-6C hi Suppressive Monocytes in Experimental Autoimmune Encephalomyelitis . J. Immunol. 179, 5228–5237 (2007).

13. V. Moliné-Velázquez, H. Cuervo, V. Vila-Del Sol, M. C. Ortega, D. Clemente, F. De Castro, Myeloid-derived suppressor cells limit the inflammation by promoting T lymphocyte apoptosis in the spinal cord of a murine model of multiple sclerosis. Brain Pathol. 21, 678–691 (2011).

14. F. Veglia, E. Sanseviero, D. I. Gabrilovich, Myeloid-derived suppressor cells in the era of increasing myeloid cell diversity. Nat. Rev. Immunol. 21 (2021), pp. 485–498.

15. L. Dolcetti, E. Peranzoni, S. Ugel, I. Marigo, A. F. Gomez, C. Mesa, M. Geilich, G. Winkels, E. Traggiai, A. Casati, F. Grassi, V. Bronte, Hierarchy of immunosuppressive strength among myeloid-derived suppressor cell subsets is determined by GM-CSF. Eur. J. Immunol. 40, 22–35 (2010).

16. C. Melero-Jerez, A. Alonso-Gómez, E. Moñivas, R. Lebrón-Galán, I. Machín-Díaz, F. de Castro, D. Clemente, The proportion of Myeloid-Derived Suppressor Cells in the spleen is related to the severity of the clinical course and tissue damage extent in a murine model of Multiple Sclerosis. Neurobiol. Dis. 140 (2020), doi:10.1016/j.nbd.2020.104869.

17. M. Mecha, A. Feliú, I. Machín, C. Cordero, F. Carrillo-Salinas, L. Mestre, G. Hernández-Torres, S. Ortega-Gutiérrez, M. L. López-Rodríguez, F. de Castro, D. Clemente, C. Guaza, 2-AG limits Theiler’s virus induced acute neuroinflammation by modulating microglia and promoting MDSCs. Glia. 66, 1447–1463 (2018).

18. C. Melero-Jerez, M. Suardíaz, R. Lebrón-Galán, C. Marín-Bañasco, B. Oliver-Martos, I. Machín-Díaz, Ó. Fernández, F. de Castro, D. Clemente, The presence and suppressive activity of myeloid-derived suppressor cells are potentiated after interferon-β treatment in a murine model of multiple sclerosis. Neurobiol. Dis. 127, 13–31 (2019).

19. V. Bronte, S. Brandau, S. H. Chen, M. P. Colombo, A. B. Frey, T. F. Greten, S. Mandruzzato, P. J. Murray, A. Ochoa, S. Ostrand-Rosenberg, P. C. Rodriguez, A. Sica, V. Umansky, R. H. Vonderheide, D. I. Gabrilovich, Recommendations for myeloid-derived suppressor cell nomenclature and characterization standards. Nat. Commun. 7 (2016), doi:10.1038/ncomms12150.

20. B. Knier, M. Hiltensperger, C. Sie, L. Aly, G. Lepennetier, T. Engleitner, G. Garg, A. Muschaweckh, M. Mitsdörffer, U. Koedel, B. Höchst, P. Knolle, M. Gunzer, B. Hemmer, R. Rad, D. Merkler, T. Korn, Myeloid-derived suppressor cells control B cell accumulation in the central nervous system during autoimmunity. Nat. Immunol. 19, 1341–1351 (2018).

21. M. Ioannou, T. Alissafi, I. Lazaridis, G. Deraos, J. Matsoukas, A. Gravanis, V. Mastorodemos, A. Plaitakis, A. Sharpe, D. Boumpas, P. Verginis, Crucial Role of Granulocytic Myeloid-Derived Suppressor Cells in the Regulation of Central Nervous System Autoimmune Disease. J. Immunol. 188, 1136–1146 (2012).

22. E. Iacobaeus, R. V. Sugars, A. Törnqvist Andrén, J. J. Alm, H. Qian, J. Frantzen, J. Newcombe, K. Alkass, H. Druid, M. Bottai, M. Röyttä, K. Le Blanc, Dynamic Changes in Brain Mesenchymal Perivascular Cells Associate with Multiple Sclerosis Disease Duration, Active Inflammation, and Demyelination. Stem Cells Transl. Med. 6, 1840–1851 (2017).

23. C. Cantoni, F. Cignarella, L. Ghezzi, B. Mikesell, B. Bollman, M. M. Berrien-Elliott, A. R. Ireland, T. A. Fehniger, G. F. Wu, L. Piccio, Mir-223 regulates the number and function of myeloid-derived suppressor cells in multiple sclerosis and experimental autoimmune encephalomyelitis. Acta Neuropathol. 133, 61–77 (2017).

24. O. Butovsky, M. P. Jedrychowski, C. S. Moore, R. Cialic, A. J. Lanser, G. Gabriely, T. Koeglsperger, B. Dake, P. M. Wu, C. E. Doykan, Z. Fanek, L. Liu, Z. Chen, J. D. Rothstein, R. M. Ransohoff, S. P. Gygi, J. P. Antel, H. L. Weiner, Identification of a unique TGF-β-dependent molecular and functional signature in microglia. Nat. Neurosci. 17, 131–143 (2014).

25. M. L. Bennett, F. C. Bennett, S. A. Liddelow, B. Ajami, J. L. Zamanian, N. B. Fernhoff, S. B. Mulinyawe, C. J. Bohlen, A. Adil, A. Tucker, I. L. Weissman, E. F. Chang, G. Li, G. A. Grant, M. G. Hayden Gephart, B. A. Barres, New tools for studying microglia in the mouse and human CNS. Proc. Natl. Acad. Sci. U. S. A. 113, E1738–E1746 (2016).

26. J. ichi Satoh, Y. Kino, N. Asahina, M. Takitani, J. Miyoshi, T. Ishida, Y. Saito, TMEM119 marks a subset of microglia in the human brain. Neuropathology. 36, 39–49 (2016).

27. T. Kuhlmann, S. Ludwin, A. Prat, J. Antel, W. Brück, H. Lassmann, An updated histological classification system for multiple sclerosis lesions. Acta Neuropathol. 133, 13–24 (2017).

28. B. Zhu, J. K. Kennedy, Y. Wang, C. Sandoval-Garcia, L. Cao, S. Xiao, C. Wu, W. Elyaman, S. J. Khoury, Plasticity of Ly-6C hi Myeloid Cells in T Cell Regulation . J. Immunol. 187, 2418–2432 (2011).

29. V. Damuzzo, L. Pinton, G. Desantis, S. Solito, I. Marigo, V. Bronte, S. Mandruzzato, Complexity and challenges in defining myeloid-derived suppressor cells. Cytom. Part B - Clin. Cytom. 88 (2015), pp. 77–91.

30. M. Novotna, M. M. Paz Soldán, N. A. Zeid, N. Kale, M. Tutuncu, D. J. Crusan, E. J. Atkinson, A. Siva, B. M. Keegan, I. Pirko, S. J. Pittock, C. F. Lucchinetti, J. H. Noseworthy, B. G. Weinshenker, M. Rodriguez, O. H. Kantarci, Poor early relapse recovery affects onset of progressive disease course in multiple sclerosis. Neurology. 85, 722–729 (2015).

31. R. Martin, M. Sospedra, M. Rosito, B. Engelhardt, Current multiple sclerosis treatments have improved our understanding of MS autoimmune pathogenesis. Eur. J. Immunol. 46 (2016), pp. 2078–2090.

32. V. Viglietta, C. Baecher-Allan, H. L. Weiner, D. A. Hafler, Loss of Functional Suppression by CD4+CD25+ Regulatory T Cells in Patients with Multiple Sclerosis. J. Exp. Med. 199, 971–979 (2004).

33. J. Haas, A. Hug, A. Viehöver, B. Fritzsching, C. S. Falk, A. Filser, T. Vetter, L. Milkova, M. Korporal, B. Fritz, B. Storch-Hagenlocher, P. H. Krammer, E. Suri-Payer, B. Wildemann, Reduced suppressive effect of CD4+CD25high regulatory T cells on the T cell immune response against myelin oligodendrocyte glycoprotein in patients with multiple sclerosis. Eur. J. Immunol. 35, 3343–3352 (2005).

34. O. W. Howell, C. A. Reeves, R. Nicholas, D. Carassiti, B. Radotra, S. M. Gentleman, B. Serafini, F. Aloisi, F. Roncaroli, R. Magliozzi, R. Reynolds, Meningeal inflammation is widespread and linked to cortical pathology in multiple sclerosis. Brain. 134, 2755–2771 (2011).

35. S. R. Choi, O. W. Howell, D. Carassiti, R. Magliozzi, D. Gveric, P. A. Muraro, R. Nicholas, F. Roncaroli, R. Reynolds, Meningeal inflammation plays a role in the pathology of primary progressive multiple sclerosis. Brain. 135, 2925–2937 (2012).

36. X. Montalban, S. L. Hauser, L. Kappos, D. L. Arnold, A. Bar-Or, G. Comi, J. de Seze, G. Giovannoni, H.-P. Hartung, B. Hemmer, F. Lublin, K. W. Rammonah, K. Selmaj, A. Traboulsee, A. Sauter, D. Masterman, P. Fontoura, S. Belachew, H. Garren, N. Mairon, P. Chin, J. S. Wolinsky, Ocrelizumab versus Placebo in Primary Progressive Multiple Sclerosis. N. Engl. J. Med. 376, 209–220 (2017).

37. S. Medina, S. Sainz de la Maza, N. Villarrubia, R. Álvarez-Lafuente, L. Costa-Frossard, R. Arroyo, E. Monreal, A. Tejeda-Velarde, E. Rodríguez-Martín, E. Roldán, J. C. Álvarez-Cermeño, L. M. Villar, Teriflunomide induces a tolerogenic bias in blood immune cells of MS patients. Ann. Clin. Transl. Neurol. 6, 355–363 (2019).

38. O. Stuve, P. Soelberg Soerensen, T. Leist, G. Giovannoni, Y. Hyvert, D. Damian, F. Dangond, U. Boschert, Effects of cladribine tablets on lymphocyte subsets in patients with multiple sclerosis: an extended analysis of surface markers. Ther. Adv. Neurol. Disord. 12 (2019), doi:10.1177/1756286419854986.

39. G. Montes Diaz, J. Fraussen, B. Van Wijmeersch, R. Hupperts, V. Somers, Dimethyl fumarate induces a persistent change in the composition of the innate and adaptive immune system in multiple sclerosis patients. Sci. Rep. 8 (2018), doi:10.1038/s41598-018-26519-w.

40. I. C. Angerer, M. Hecker, D. Koczan, L. Roch, J. Friess, A. Rüge, B. Fitzner, N. Boxberger, I. Schröder, K. Flechtner, H. J. Thiesen, A. Winkelmann, S. Meister, U. K. Zettl, Transcriptome profiling of peripheral blood immune cell populations in multiple sclerosis patients before and during treatment with a sphingosine-1-phosphate receptor modulator. CNS Neurosci. Ther. 24, 193–201 (2018).

41. M. Kaufmann, R. Haase, U. Proschmann, T. Ziemssen, K. Akgün, Real-World Lab Data in Natalizumab Treated Multiple Sclerosis Patients Up to 6 Years Long-Term Follow Up. Front. Neurol. 9 (2018), doi:10.3389/fneur.2018.01071.

42. Z. Wang, G. Zheng, G. Li, M. Wang, Z. Ma, H. Li, X. Y. Wang, H. Yi, Methylprednisolone alleviates multiple sclerosis by expanding myeloid-derived suppressor cells via glucocorticoid receptor β and S100A8/9 up-regulation. J. Cell. Mol. Med. 24, 13703–13714 (2020).

43. I. L. King, T. L. Dickendesher, B. M. Segal, Circulating Ly-6C + myeloid precursors migrate to the CNS and play a pathogenic role during autoimmune demyelinating disease. Blood. 113, 3190–3197 (2009).

44. M. A. Ingersoll, A. M. Platt, S. Potteaux, G. J. Randolph, Monocyte trafficking in acute and chronic inflammation. Trends Immunol. 32 (2011), pp. 470–477.

45. G. Locatelli, D. Theodorou, A. Kendirli, M. J. C. Jordão, O. Staszewski, K. Phulphagar, L. Cantuti-Castelvetri, A. Dagkalis, A. Bessis, M. Simons, F. Meissner, M. Prinz, M. Kerschensteiner, Mononuclear phagocytes locally specify and adapt their phenotype in a multiple sclerosis model. Nat. Neurosci. 21, 1196–1208 (2018).

46. D. A. Giles, J. M. Washnock-Schmid, P. C. Duncker, S. Dahlawi, G. Ponath, D. Pitt, B. M. Segal, Myeloid cell plasticity in the evolution of central nervous system autoimmunity. Ann. Neurol. 83, 131–141 (2018).

47. V. E. Miron, A. Boyd, J. W. Zhao, T. J. Yuen, J. M. Ruckh, J. L. Shadrach, P. Van Wijngaarden, A. J. Wagers, A. Williams, R. J. M. Franklin, C. Ffrench-Constant, M2 microglia and macrophages drive oligodendrocyte differentiation during CNS remyelination. Nat. Neurosci. 16, 1211–1218 (2013).

48. Z. Yu, D. Sun, J. Feng, W. Tan, X. Fang, M. Zhao, X. Zhao, Y. Pu, A. Huang, Z. Xiang, L. Cao, C. He, MSX3 switches microglia polarization and protects from inflammation-induced demyelination. J. Neurosci. 35, 6350–6365 (2015).

49. G. Rizzo, R. Di Maggio, A. Benedetti, J. Morroni, M. Bouche, B. Lozanoska-Ochser, Splenic Ly6Chi monocytes are critical players in dystrophic muscle injury and repair. JCI Insight. 5 (2020), doi:10.1172/jci.insight.130807.

50. C. Melero-Jerez, B. Fernández-Gómez, R. Lebrón-Galán, M. C. Ortega, I. Sánchez-de Lara, A. C. Ojalvo, D. Clemente, F. de Castro, Myeloid-derived suppressor cells support remyelination in a murine model of multiple sclerosis by promoting oligodendrocyte precursor cell survival, proliferation, and differentiation. Glia. 69, 905–924 (2021).

51. F. Mei, K. Lehmann-Horn, Y. A. A. Shen, K. A. Rankin, K. J. Stebbins, D. S. Lorrain, K. Pekarek, S. A. Sagan, L. Xiao, C. Teuscher, H. C. von Büdingen, J. Wess, J. Josh Lawrence, A. J. Green, S. P. J. Fancy, S. S. Zamvil, J. R. Chan, Accelerated remyelination during inflammatory demyelination prevents axonal loss and improves functional recovery. Elife. 5 (2016), doi:10.7554/eLife.18246.

52. V. A. Deshmukh, V. Tardif, C. A. Lyssiotis, C. C. Green, B. Kerman, H. J. Kim, K. Padmanabhan, J. G. Swoboda, I. Ahmad, T. Kondo, F. H. Gage, A. N. Theofilopoulos, B. R. Lawson, P. G. Schultz, L. L. Lairson, A regenerative approach to the treatment of multiple sclerosis. Nature. 502, 327–332 (2013).

53. I. D. Duncan, A. Brower, Y. Kondo, J. F. Curlee, R. D. Schultz, Extensive remyelination of the CNS leads to functional recovery. Proc. Natl. Acad. Sci. U. S. A. 106, 6832–6836 (2009).

54. R. M. Bove, A. J. Green, Remyelinating Pharmacotherapies in Multiple Sclerosis. Neurotherapeutics. 14 (2017), pp. 894–904.

55. B. Bodini, M. Veronese, D. García-Lorenzo, M. Battaglini, E. Poirion, A. Chardain, L. Freeman, C. Louapre, M. Tchikviladze, C. Papeix, F. Dollé, B. Zalc, C. Lubetzki, M. Bottlaender, F. Turkheimer, B. Stankoff, Dynamic Imaging of Individual Remyelination Profiles in Multiple Sclerosis. Ann. Neurol. 79, 726–738 (2016).

56. C. Lubetzki, B. Zalc, A. Williams, C. Stadelmann, B. Stankoff, Remyelination in multiple sclerosis: from basic science to clinical translation. Lancet Neurol. 19 (2020), pp. 678–688.

57. J.-I. Youn, S. Nagaraj, M. Collazo, D. I. Gabrilovich, Subsets of Myeloid-Derived Suppressor Cells in Tumor-Bearing Mice. J. Immunol. 181, 5791–5802 (2008).

58. D. I. Gabrilovich, S. Nagaraj, Myeloid-derived suppressor cells as regulators of the immune system. Nat. Rev. Immunol. 9 (2009), pp. 162–174.

59. P. Y. Pan, G. Ma, K. J. Weber, J. Ozao-Choy, G. Wang, B. Yin, C. M. Divino, S. H. Chen, Immune stimulatory receptor CD40 is required for T-cell suppression and T regulatory cell activation mediated by myeloid-derived suppressor cells in cancer. Cancer Res. 70, 99–108 (2010).

60. V. Moliné-Velázquez, M. C. Ortega, V. Vila del Sol, C. Melero-Jerez, F. de Castro, D. Clemente, The synthetic retinoid Am80 delays recovery in a model of multiple sclerosis by modulating myeloid-derived suppressor cell fate and viability. Neurobiol. Dis. 67, 149–164 (2014).

61. Y. Dombrowski, T. O’Hagan, M. DIttmer, R. Penalva, S. R. Mayoral, P. Bankhead, S. Fleville, G. Eleftheriadis, C. Zhao, M. Naughton, R. Hassan, J. Moffat, J. Falconer, A. Boyd, P. Hamilton, I. V. Allen, A. Kissenpfennig, P. N. Moynagh, E. Evergren, B. Perbal, A. C. Williams, R. J. Ingram, J. R. Chan, R. J. M. Franklin, D. C. Fitzgerald, Regulatory T cells promote myelin regeneration in the central nervous system. Nat. Neurosci. 20, 674–680 (2017).

62. S. Bramow, J. M. Frischer, H. Lassmann, N. Koch-Henriksen, C. F. Lucchinetti, P. S. Sørensen, H. Laursen, Demyelination versus remyelination in progressive multiple sclerosis. Brain. 133, 2983–2998 (2010).

63. E. C. W. Breij, B. P. Brink, R. Veerhuis, C. Van Den Berg, R. Vloet, R. Yan, C. D. Dijkstra, P. Van Der Valk, L. Bö, Homogeneity of active demyelinating lesions in established multiple sclerosis. Ann. Neurol. 63, 16–25 (2008).

64. A. F. Lloyd, V. E. Miron, The pro-remyelination properties of microglia in the central nervous system. Nat. Rev. Neurol. 15 (2019), pp. 447–458.

65. D. Clemente, M. C. Ortega, F. J. Arenzana, F. de Castro, FGF-2 and anosmin-1 are selectively expressed in different types of multiple sclerosis lesions. J. Neurosci. 31, 14899–14909 (2011).

66. L. A. Boven, M. Van Meurs, M. Van Zwam, A. Wierenga-Wolf, R. Q. Hintzen, R. G. Boot, J. M. Aerts, S. Amor, E. E. Nieuwenhuis, J. D. Laman, Myelin-laden macrophages are anti-inflammatory, consistent with foam cells in multiple sclerosis. Brain. 129, 517–526 (2006).

67. N. L. Fransen, C. C. Hsiao, M. Van Der Poel, H. J. Engelenburg, K. Verdaasdonk, M. C. J. Vincenten, E. B. M. Remmerswaal, T. Kuhlmann, M. R. J. Mason, J. Hamann, J. Smolders, I. Huitinga, Tissue-resident memory T cells invade the brain parenchyma in multiple sclerosis white matter lesions. Brain. 143, 1714–1730 (2020).

68. A. C. Ruifrok, D. A. Johnston, Quantification of histochemical staining by color deconvolution. Anal. Quant. Cytol. Histol. 23, 291–299 (2001).

