## Supplementary material for "Myeloid-Derived Suppressor Cells are relevant factors to predict the severity of multiple sclerosis"

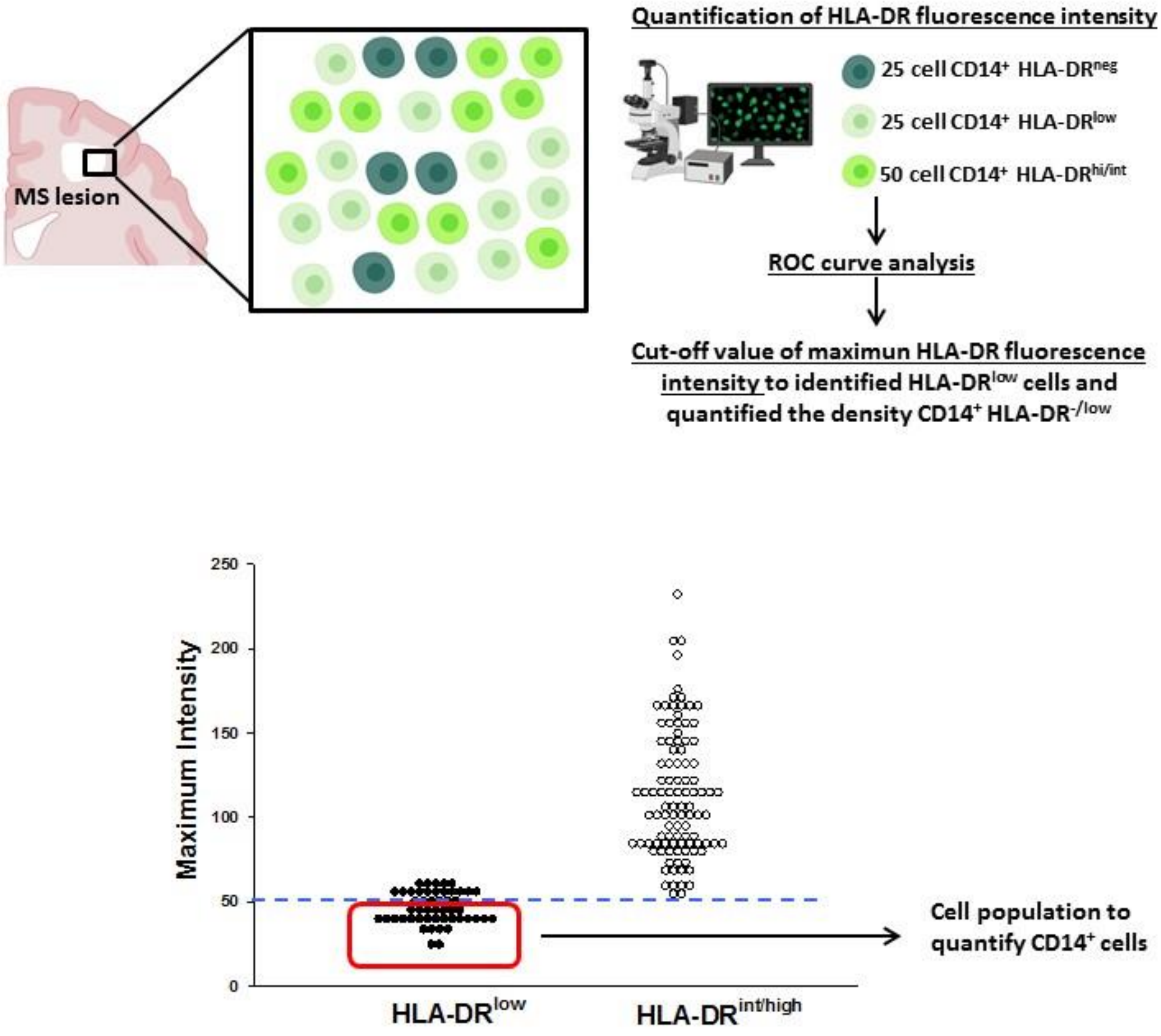


**Fig. S1. A**: Scheme of the HLA-DR cell diversity in MS lesions according to their fluorescence intensity. **B**: Representative quantification of HLA-DR intensity in a MS patient in order to identify the HLA-DR$^{low}$ cell population based on the cut-off value.

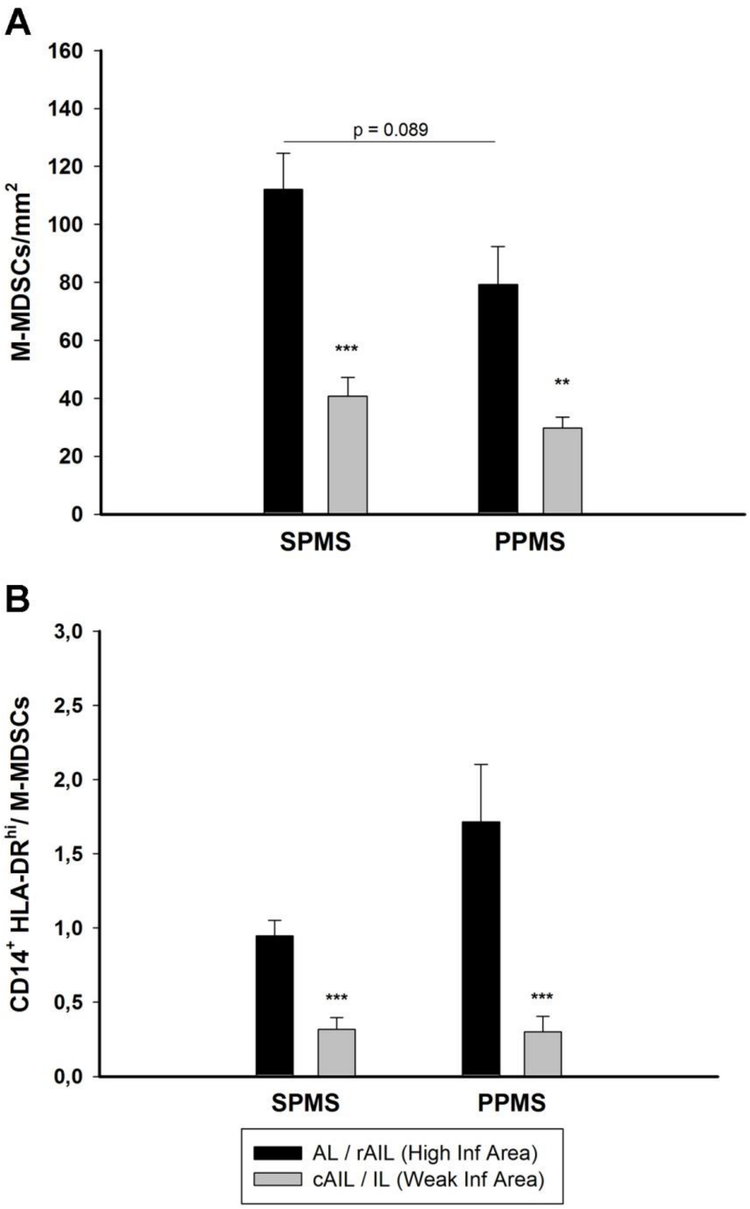


**Fig. S2. A:** Density of M-MDSCs in high inflammatory areas (AL and rAIL) in SPMS and PPMS patients was similar and significantly higher than in weak inflammatory areas (cAIL and IL, 17 SPMS and 12 PPMS patients). **B**: Similar to M-MDSCs, the IIR (balance between pro-inflammatory $CD14^+$ $HLA\text{-}DR^{hi}$ cells and M-MDSCs) was significantly higher in high inflammatory regions of both SPMS and PPMS. The comparative analysis was carried out with a Student's t test and the mean (± SEM) is shown: ** $p < 0.01$ and ***$p < 0.001$.

| | High Inflammatory area | | | | Weak Inflammatory area | | | |
|---|---|---|---|---|---|---|---|---|
| | *SPMS* | | *PPMS* | | *SPMS* | | *PPMS* | |
| | M-MDSCs | IIR | M-MDSCs | IIR | M-MDSCs | IIR | M-MDSCs | IIR |
| **TP (h)** | r = -0.192<br>p = 0.530 | r = 0.388<br>p = 0.19 | r = -0.125<br>p = 0.730 | r = 0.486<br>p =0.137 | r = -0.092<br>p = 0.928 | r = -0.0845<br>p =0.783 | r = -0.065<br>p = 0.057 | r = -0.141<br>p = 0.681 |
| **Age (y)** | r = -0.394<br>p = 0.131 | r = 0.263<br>p = 0.326 | **r = 0.613**<br>**p = 0.045** | r = -0.266<br>p =0.416 | r = -0.288<br>p = 0.318 | r = 0.006<br>p = 0.976 | r = 0.078<br>p = 0.830 | r = -0.248<br>p = 0.450 |

**Table S1:** Density of M-MDSCs and inflammatory-immunoregulatory ratio ($CD14^+$ HLA-$DR^{hi}$/M-MDSC). *Abbreviations:* h, hours; IIR, inflammatory-immunoregulatory ratio; PPMS, primary progressive multiple sclerosis; SPMS: secondary progressive multiple sclerosis; TP, time postmortem; y, years.

| Epidemiological data | All MS patients n = 39 | MS ≤30 days n = 23 | MS >30 days n = 16 | HC n = 26 |
|---|---|---|---|---|
| **Age**[a] | 37.39 (9.81) | 36.09 (7.87) | 39.25 (8.61) | 34.77 (10.28) |
| **Male/female (%female)** | 13/26 (66.67) | 9/14 (60.87) | 4/12 (75.00) | 12/14 (53.84) |
| **EDSS baseline**[b] | 3 (2-4) | 3 (3-4) | 2 (0.5-2) | NA |
| **EDSS 12m**[b] | 0 (0-2) | 0 (0-1) | 1 (0-2) | NA |

**Table S2**. Demographic table of the MS patients enrolled for the PMBC follow up study. *Abbreviations:* HC: Healthy controls. [a] = mean ± SD. [b] = median ± interquartile range. NA: Not applicable.

| Patient ID | I | II | III | IV | V | VI | VII | VIII | IX | X | XI | XII | XIII | XIV | XV | XVI | XVII | XVIII | XIX | XX | XXI | XXII | XXIII |
|---|---|---|---|---|---|---|---|---|---|---|---|---|---|---|---|---|---|---|---|---|---|---|---|
| **Age** | 44 | 42 | 51 | 24 | 30 | 50 | 32 | 42 | 40 | 44 | 30 | 42 | 39 | 23 | 57 | 37 | 29 | 37 | 32 | 30 | 36 | 16 | 24 |
| **Sex** | F | M | M | F | M | F | F | F | F | F | F | M | F | M | F | M | M | M | M | F | F | F | M |
| **Time since relapse (days)** | 10 | 8 | 22 | 6 | 5 | 21 | 33 | 10 | 32 | 19 | 12 | 21 | 6 | 11 | 16 | 17 | 13 | 14 | 16 | 30 | 12 | 5 | 9 |
| **Treatment during follow-up** | ---- | R | DF | Cpx | Cc DF | Cc DF | Cc Hf | Cc Hf Cpx | Cc DF | Cc Cdb | Cc DF | Cc | Hf | Cc DF | Cc | DF | Cc DF Hf | Cc DF | Cc DF | Cc PG | Cc DF | Ntz | DF |
| **EDSS baseline** | 2 | 2 | 4 | 3 | 3 | 3.5 | 4 | 5 | 4 | 4 | 4 | 3 | 4 | 4 | 3 | 2 | 3 | 3 | 2 | 3 | 2 | 3 | 2 |
| **EDSS 12months** | 0 | 0 | 1 | 0 | 0 | 3 | 0 | 4 | 3 | 0 | 1 | 2 | 1 | 0 | 0 | 0 | 0 | 0 | 0 | 1 | 0 | 2 | 0 |
| **MDSCs/MNCs baseline** | 2.67 | 2.27 | 0.52 | 2.00 | 0.82 | 1.41 | 2.32 | 0.7 | 0.07 | 0.72 | 0.36 | 0.74 | 1.38 | 0.15 | 0.76 | 1.41 | 2.46 | 0.06 | 1.37 | 1.11 | 1.85 | 0.30 | 3.39 |
| **MDSCs/MNCs 12months** | 1.62 | 0.92 | 0.87 | 3.47 | 1.69 | 4.05 | 3.14 | 0.28 | 7.34 | 0.89 | 0.88 | 2.79 | 1.48 | 1.11 | 1.59 | 7.71 | 0.31 | 2.54 | 5.26 | 1.44 | 6.53 | 1.09 | 0.95 |

**Table S3**. Clinical and demographic data of MS followed up patients. *Abbreviations:* Cc: corticosteroids; Cdb: Cladribina; Cpx: Copaxone; DF: Dimethyl Fumarate; Hf: Hidroferol; R: Rebif 44; Ntz: Natalizumab; PG: Plegridy
